# Systematic microcircuit reconfigurations underlie early experience-induced visual cortical plasticity

**DOI:** 10.64898/2025.12.21.695847

**Authors:** Li Yao, Ziwei Shang, Zilong Ji, Meizhen Meng, Fengchao Wang, Xiang Gu, Miao He, Si Wu, Xiaohui Zhang

**Affiliations:** State Key Laboratory of Cognitive Neuroscience & Learning, IDG/McGovern Institute for Brain Research, Beijing Normal University, Beijing 100875, China; School of Psychology and Cognitive Sciences, IDG/McGovern Institute for Brain Research, Center of Quantitative Biology, Peking-Tsinghua Center for Life Sciences, Peking University, Beijing 100875, China; Institutes of Brain Science, Fudan University, Shanghai 200032, China

**Author notes:** These authors contributed equally (Li Yao, Ziwei Shang, Zilong Ji).

**Keywords:** Primary visual cortex, Experience-dependent plasticity, Disinhibitory synapse, Microcircuit

## Abstract

Experience-dependent reorganization of neuronal connections during a postnatal critical period (CP) is essential for the functional maturation of developing primary visual cortex (V1). However, the systematic reconfigurations across microcircuit synapses underlying this plasticity were poorly defined. Paired recording from excitatory pyramidal cells and other three major interneurons (INs) within layers 2/3 and 4 microcircuits in the developing mouse V1 monocular region, we find that monocular deprivation (MD) for one-day during the CP transiently reconfigures local excitatory synapses on inhibitory INs that specifically express parvalbumin (PV) or vasoactive intestinal peptide (VIP), whereas prolonged MD primarily strengthens inhibitory transmission from somatostatin (SST)-expressing INs to PV-INs and VIP-INs in both layers. A physiologically based microcircuit model suggests these coordinated synaptic changes in specific inhibitory neurons mediate differential MD-induced plasticity in visual responses across layer 2/3 neurons *in vivo*. Together, our findings define a systematic microcircuit basis for cortical critical-period plasticity.

**Highlights:**

- Inhibitory-to-inhibitory connectivity is conserved in cortical layer 2/3 and 4 circuits.
- Early experience sequentially modifies excitatory and dis-inhibitory synapses in both layers.
- Systematic microcircuit reconfigurations underlie plasticity of cortical visual responses.
- Modeling suggests a pivotal role of disinhibitory microcircuits in mediating the plasticity.

## Introduction

Neuronal connections in developing sensory and higher-function cortices undergo extensive experience-dependent refinements during a postnatal critical period (CP). In the developing sensory cortex, this heightened plasticity is essential for the maturation of cortical sensory processing functions^1–4^. A classical model for studying this plasticity process is the ocular dominance plasticity (ODP) of developing mammalian primary visual cortex (V1), characterized by profound and permanent shifts in the preference of visual spiking responses of V1 neurons following monocular deprivation (MD) during the CP^5–8^. Although the cortex is organized into six layers^9^, it is generally known that MD-induced ODP is predominantly expressed in layer 2/3 and 4 of developing V1^10,11^ Accumulative evidence suggested that CP experience modifies both excitatory and inhibitory synapses within cortical microcircuits in a sequential manner^12–14^. For example, brief MD (1-day) transiently reduced excitatory synaptic drive onto specific subtypes of inhibitory interneurons (INs) in layer 2/3, while prolonged MD (3-5 days) enhanced GABAergic transmission on pyramidal cells (PCs)^15,16^. Moreover, recent studies implicated disinhibitory circuits - synapses between cortical inhibitory interneurons (INs) might be involved in the process of regulating the initiation and closure of the CP^17,18^. These findings indicate that MD-induced circuit reorganization arises from dynamic interactions between excitatory and inhibitory microcircuits across different layers and over time^8,19,20^. However, how these microcircuits are reconfigured spatiotemporally remains a fundamental unanswered question^4^, and elucidating this systematic reconfiguration is critical for uncovering the synaptic mechanisms underlying experience-dependent cortical plasticity.

Cortical layers comprise dominant excitatory PCs and diverse subtypes of inhibitory INs-including those expressing parvalbumin (PV), somatostatin (SST), or vasoactive intestinal peptide (VIP). These cellular elements form relatively canonical intra-laminar synaptic microcircuits. For example, PV-INs primarily establish strong connections with neighboring PCs and other PV cells; SST-INs inhibit most other cortical neurons but exhibit self-avoidance, and VIP-INs preferentially target SST-INs^21,22^. The layer 2/3 microcircuit possesses the highest degree of recurrent synaptic connectivity among PCs and various INs subtypes^23^. However, emerging evidence from electrophysiological recordings and volume electron-microscopy (EM) reconstruction has revealed greater microcircuit diversity^24–27^ and non-canonical inhibitory connection motifs across cortical layers than previously recognized^28^. These layer-specific synaptic architectures likely play dissociable roles in regulating cortical computations^29,30^ and experience-dependent plasticity^20^. Although the ODP is expressed most prominently in both layers 2/3 and 4, it is mediated by distinct synaptic and molecular mechanisms in each^11,31^. Thus, identifying the specific loci of experience-induced synaptic modifications across cortical layers is essential for a systematical dissection of the circuit mechanism underlying cortical plasticity.

In the present study, we conducted extensive *ex vivo* and *in vivo* recordings from PCs, PV-, SST-and VIP-INs within the monocular region of developing mouse V1 (V1M) following 1-day or 4-day MD at the CP peak time. We identified specific loci of MD-induced synaptic changes in intra- or inter-laminar microcircuits across layers 2/3 and 4, respectively, over time, and found that predominant synaptic reconfigurations reside at those synapses either on INs or between INs in both layers. Further computations using a physiologically grounded microcircuit model suggested these coordinated synaptic changes in specific inhibitory neurons mediate differential MD-induced plasticity in visual responses of layer-2/3 neurons *in vivo*. Thus, our results provide a systematic microcircuit basis for cortical experience-dependent plasticity.

## Results

All *ex vivo* or *in vivo* electrophysiological recordings were performed across layers 2/3–4 within the V1M of mice aged postnatal days 27–32 (P27–32), a time window corresponding to the peak of OD for mouse ODP^2^. The V1M receives inputs exclusively from the contralateral eye and lacks the process of binocular synaptic competition, thereby as an ideal region for the specific characterization of MD-induced synaptic circuit changes^8,19^. In experiments, we used the following IN subtype-specific reporter mouse lines, *PV-cre::Ai9* (tdTomato), *SST-cre::Ai9* and *VIP-cre::Ai9*, to record specifically from cortical PV-, SST-, and VIP-INs, respectively. Each line was further crossed with the *B13*^32^ or *GIN*^33^ transgenic BAC lines, which enabled simultaneous genetic identification of two IN-subtypes within the V1M through the expression of different fluorescence proteins.

### Distinct and conserved synaptic wirings between layer 2/3 and 4 microcircuits in the V1M

We first performed whole-cell recordings from paired cortical neurons (spaced within 50 μm horizontally or vertically) in the V1M of acutely prepared visual cortical slices from norm-reared mice aged P27–28. Recordings targeted excitatory PCs and inhibitory PV-, SST- and VIP-INs in layers 2/3 and 4, respectively. A total of 3,585 chemical synapses were tested by delivering a train of five spikes at 20 Hz to the presynaptic neuron (current-clamp mode) to evoke either unitary excitatory or inhibitory postsynaptic currents (uEPSCs or uIPSCs) in connected postsynaptic neurons (**Figure 1A** and **S1**).

**Figure 1.**
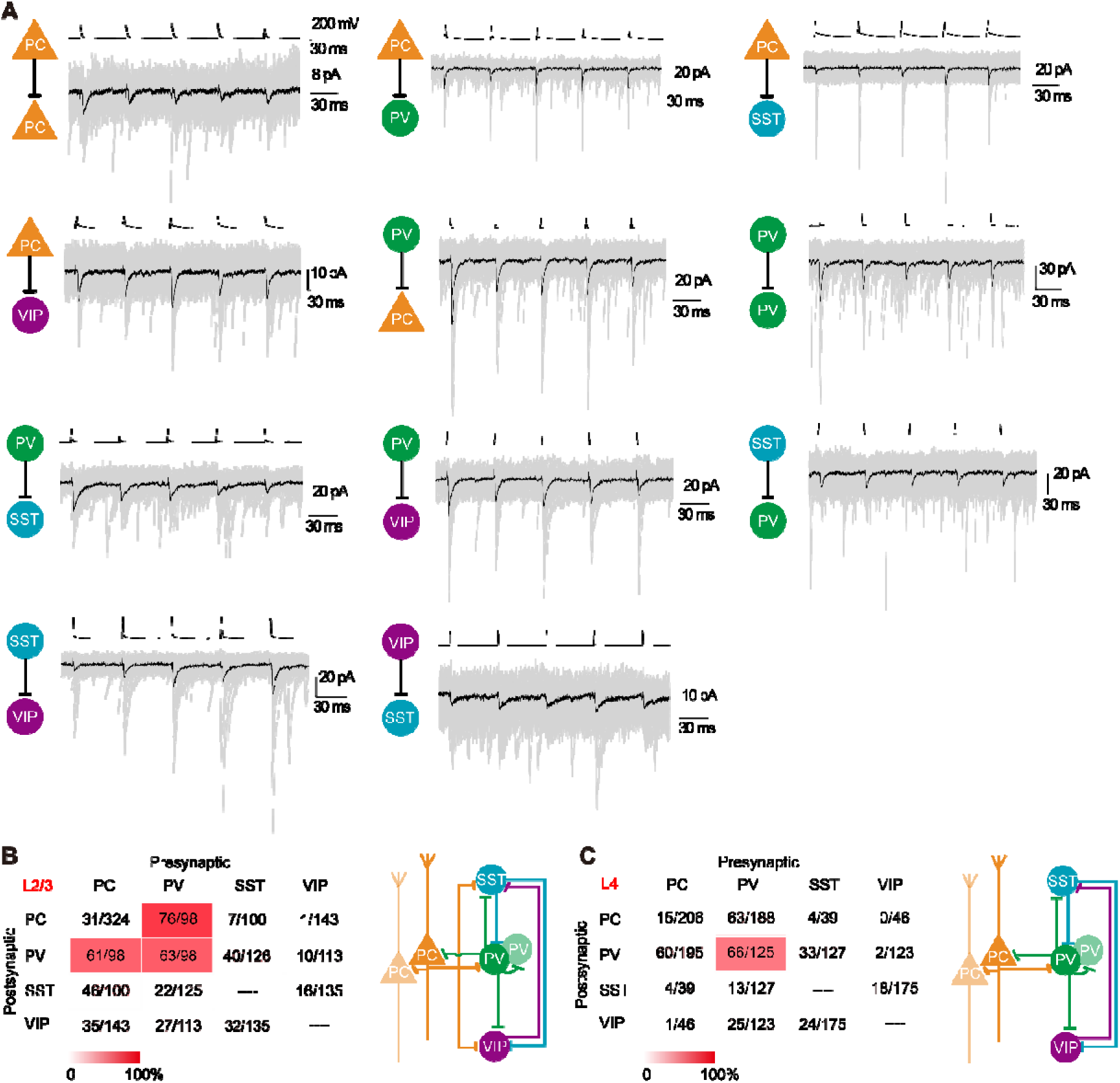
Shared and distinct synaptic organizations in the layer 2/3 and 4 microcircuits of developing visual cortex. (A) Representative traces of uEPSCs or uIPSCs recorded from various excitatory, inhibitory or disinhibitory synapses among PCs, PV-, SST- and VIP-INs in the layer 2/3 microcircuit. Synaptic transmissions were tested by 5 presynaptic spikes at 20Hz for 30 trials. Gray and black lines represent evoked raw traces and average traces, respectively. (B) A matrix of connection probability among the four types of neurons in V1 layer 2/3 (left). Schematic diagram of the layer 2/3 microcircuit organization (right). (C) A matrix of connection probability among the four types of neurons in V1 layer 4 (left). Schematic diagram of the layer 4 microcircuit organization (right).

By quantifying the probability of tested synaptic connections, we generated connection maps of layer 2/3 and 4 microcircuits (**Figures 1B** and **1C**). Excitatory synapses from PCs to neighboring PV-, SST- or VIP-INs exhibited significantly higher probabilities in layer 2/3, compared to layer 4 (PC-PV, 62.24% and 30.77% in layer 2/3 and 4, *p* < 1 × 10^-4^; PC-SST, 46% and 10.26%, *p* < 1× 10^-4^, Chi-square test) and in the layer 4 PC synapses barely targeted VIP-INs (PC-VIP, 24.48% and 2.17%, *p* = 8 ×10^-4^, Chi-square test; **Figures 1B** and **1C**, **Table S1**). In contrast, the probability of PC-PC excitatory connections did not differ between layers. Moreover, for inhibitory-to-excitatory (I-E) microcircuits within a same layer, PV-INs formed inhibitory synapses on PCs at a higher probability, while SST-INs did so at a lower probability (layer 2/3: PV-PC, 77.55%, SST-PC 7%, *p* < 1 × 10^-4^; layer 4: 33.51% and 10.26%, *p* = 0.0038, Chi-square test). VIP-INs rarely form inhibitory synapses on PCs in either layer (**Table S1**). Notably, in layer 2/3, PV-INs connected to approximately 78% neighboring PCs, a value substantially higher than layer 4 (PV-PC, 77.55% and 33.51% in layer 2/3 and 4, *p* < 1 × 10^-4^). However, the unitary strength at all tested synapses formed between PCs and INs did not differ between layers (**Table S1**). These findings suggested that layers 2/3 and 4 possess distinct synaptic microcircuits between excitatory and inhibitory neurons in developing V1 during the CP.

In contrast, canonical patterns of inhibitory-to inhibitory (IN-IN) synaptic microcircuits were conserved across layers. As shown by the **Figures 1B, 1C** and **Table S1**, the connection probabilities of tested IN-IN inhibitory synapses (also referred to as disinhibitory synapses in the context of PCs) were highly similar between layers, with the exception of SST-VIP (SST-VIP, 23.70 % and 13.71% in layers 2/3 and 4, respectively, *p* = 0.023, Chi-square test) and VIP-PV (VIP-PV, 8.85 % and 1.63% in layers 2/3 and 4, respectively, *p* = 0.012) disinhibitory connections that exhibited a relatively higher probability in layer 2/3. While unitary strengths of these disinhibitory synapses were comparable between layers, the short-term dynamics of disinhibitory synaptic transmission involving VIP-INs differed substantially. For example, PV-VIP disinhibitory transmission in both layers showed short-term depression, with greater depression observed in layer 4. Conversely, SST-VIP and VIP-SST disinhibitory synapses showed short-term facilitation, with greater facilitation in layers 2/3 and 4, respectively (**Figure 1A**, **Figures S2** and **S3**). Moreover, In agree with previous studies^34,35^, our result also indicate that the short-term dynamics of cortical excitatory transmission depend on the subtypes of postsynaptic neurons, whereas the dynamics of inhibitory and disinhibitory transmission are primarily determined by the subtypes of presynaptic INs subtypes.

Collectively, these results highlight both distinct and conserved features of synaptic microcircuits in layer 2/3 and 4 of developing mouse V1. These laminar distinct features in cortical microcircuit connectivity were also observed in the matured V1^25,27^.

### 1-day MD selectively reconfigures local excitatory synapses in layers 2/3 and 4

We next systematically measured the MD effects on developing local excitatory synapses in layers 2/3 or 4. Mice underwent MD for 1 or 4 days starting at P27 or P28, with age-matched normally-reared mice severing as controls.

We first examined recurrent excitatory synapses targeting either neighboring PCs and three IN subtypes, respectively, within each layer (**Figures 2A** and **2F**). We found, in layer 2/3, both 1-day and 4-day MD did not alter the amplitude of uEPSC of PC-PC excitatory synapses (*p* = 0.35 and *p* = 0.48, respectively, Mann-Whitney U test; Figure 2B). In contrast, 1-day MD selectively enhanced the strength of PC-PV and PC-VIP excitatory synapses in layer 2/3 (*p* = 0.029 and *p* = 0.029; **Figures 2C** and **2E**), but not at PC-SST excitatory synapses (*p* = 0.65; **Figure 2D**). Notably, these enhancements were transient because they were no longer present after 4-day MD (*p* = 0.99 and *p* = 0.75 for PC-PV and PC-VIP, respectively; **Figures 2C** and **2E**).

**Figure 2.**
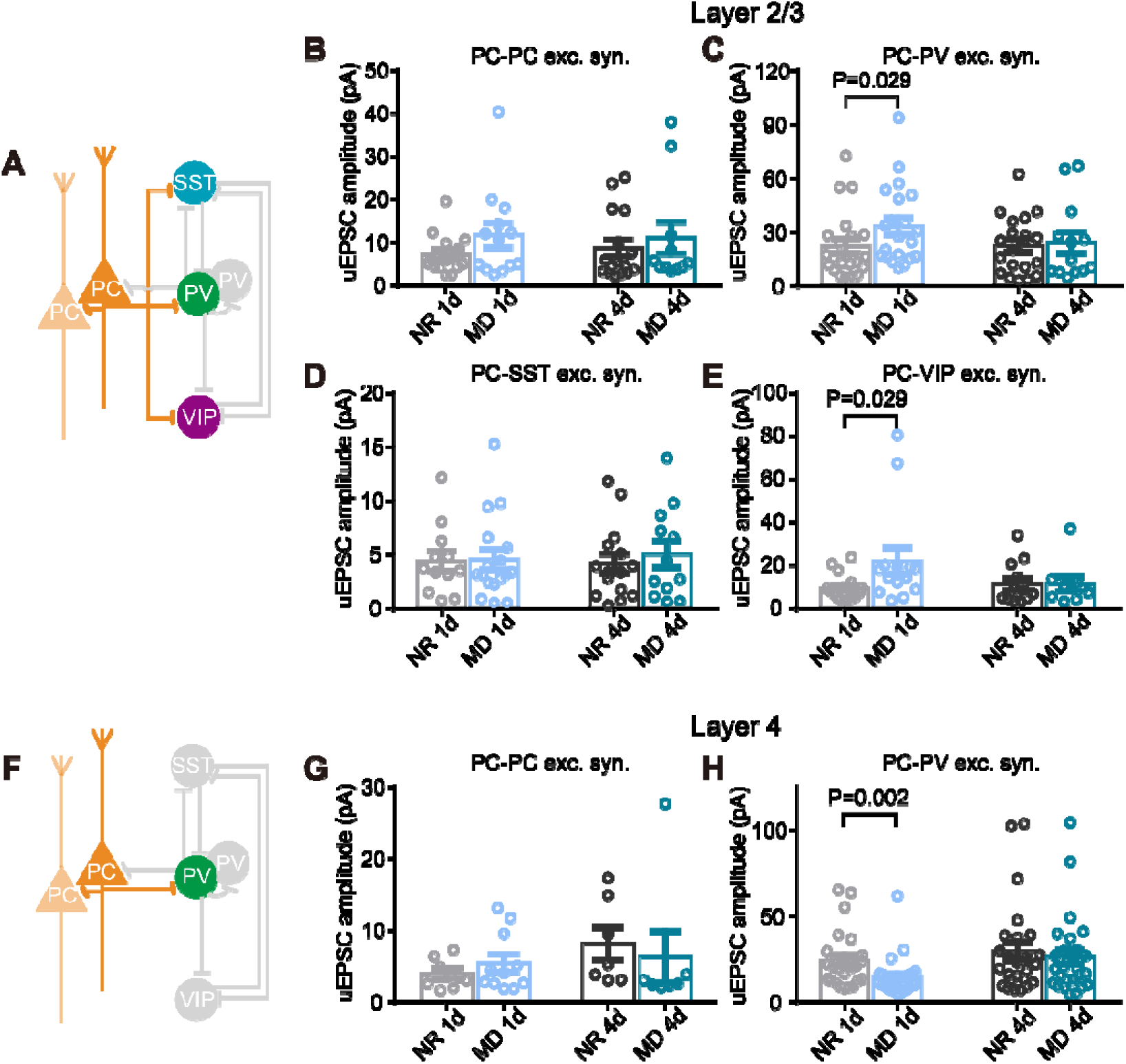
Selective modification of excitatory synapses on INs in both layers by 1-day MD during the CP. (A) A diagram illustrating examined excitatory synapses (in colors) within the layer 2/3 microcircuit. (B– E) Changes of synaptic strengths of excitatory synapse on PCs (B), PV-INs (C), SST-Ins (D) or VIP-INs (E) in the layer 2/3, following 1 or 4-day MD and age-matched normal rearing (NR 1d and NR 4d) during the critical period, respectively. (F) A diagram illustrating examined excitatory synapses (in colors) within the layer 4 microcircuit. (G, H) Changes of synaptic strengths of excitatory synapse on PCs (g), PV-INs (h) in the layer 4, following MD for 1 day (MD 1d) or 4 days (MD 4d) and age-matched normal rearing (NR 1d and NR 4d) during the critical period, respectively. Data are presented in mean ± s.e.m., the dots represent the connected synapses. See statistical data in Table S2.

A similar assessment in layer 4 indicated that 1- or 4-day MD did not affect PC-PC synapse as well (*p* = 0.60 and *p* = 0.07, respectively, Mann-Whitney *U* test; **Figure 2G**). However, 1-day MD reduced the efficacy of PC-PV excitatory synapses in this layer (*p* = 0.002; **Figure 2H**). Due to their low connection probability in layer 4, we did not examine PC-SST or PC-VIP synapses for all conditions. Taken together, these results demonstrate that 1-day MD induces a rapid and selective reconfigurations of local excitatory transmission onto specific IN subtypes in a layer-specific manner: PC-PV and PC-VIP excitatory synapses are enhanced in layer 2/3, while PC-PV synapses are weakened in layer 4. Notably, all these 1-day MD-induced synaptic modifications are transient, returning to baseline after 4 days of MD.

### 4-day MD selectively reduced local PV-synapses and layer 4 excitatory inputs on PCs in layer 2/3

We next examined how MD alters local inhibitory synapses on PCs within layer 2/3 and 4, respectively. Previous studies^21,36^ and our current results (**Figure 1**) indicate that PV-INs could mediate major and stronger feed-forward or lateral inhibition by innervating nearby PCs with generally higher probability within both layers, compared to SST-INs and VIP-INs. Our first experiment thus focused on PV-PC inhibitory synapses. We found that in layer 2/3, 1-day MD did not alter the amplitude of uIPSCs at PV-PC synapses (*p* = 0.49, Mann-Whitney *U* test; **Figure 3A**). However, 4-day MD robustly decreased uIPSC amplitudes (*p* = 0.005; **Figure 3A**). Notably, 1-day MD increased the connection probability of these synapses in layer 2/3 (*p* = 0.044, Chi-square test; **Figure S3E**), an effect that was not sustained after 4 days of MD (*p* = 0.24, Mann-Whitney *U* test; **Figure S3E**). This transient remodeling of synaptic connectivity is consistent with a recent study^15^. In contrast to layer 2/3, neither 1-day nor 4-day MD affected uIPSC amplitudes at PV-PC synapses in layer 4 (*p* = 0.26 and *p* = 0.17, respectively; **Figure 3B**). These results indicate that MD-induced suppression of inhibitory synaptic transmission selectively recruits PV-PC synapses in layer 2/3 only by 4-day MD.

**Figure 3.**
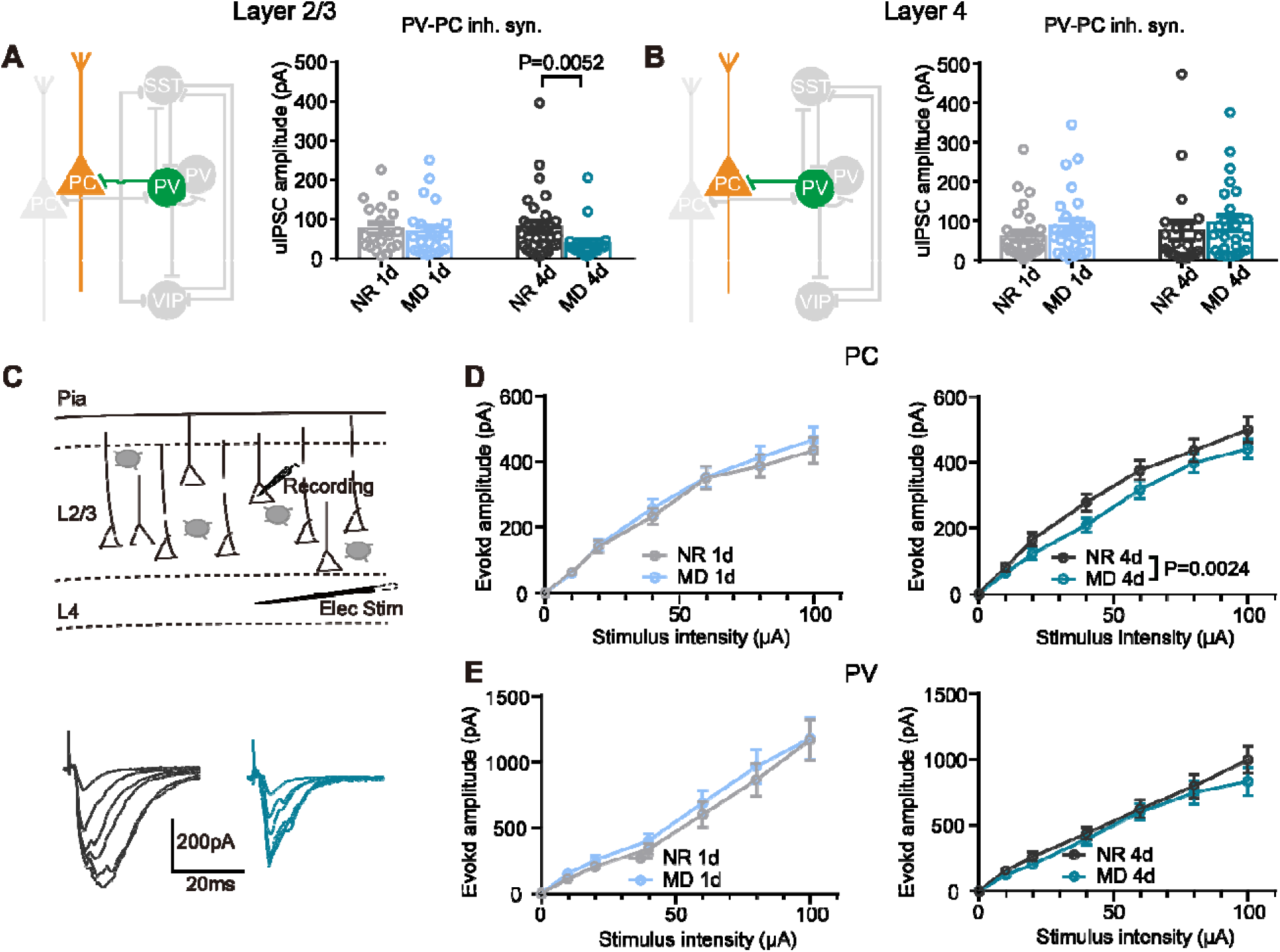
4-day MD selectively decreases the layer 2/3 PV-PC inhibitory and the translaminar inputs from layer 4. (A, B) Effects of MD 1d and 4d on the PV-PC inhibitory synapses in both layer 2/3 (A) and layer 4 (B) microcircuits. Diagrams showing the examined synapse (in colors). (C) A diagram illustrating experimental configuration for recording EPSCs in layer 2/3 PC in response to a field electrical stimulation to layer 4 PCs in the V1M slice (top). Representativ evoked EPSCs of layer 2/3 PC with increased stimulation intensities in the age-matched NR 4d (black) and MD 4d (cyan) mice, respectively (bottom). (D) Summarized changes of the layer 4-to-layer 2/3 PCs translaminar synaptic strength following the MD 1d (left) and MD 4d (right), respectively. (E) Summarized changes of the layer 4-to-layer 2/3 PV-INs translaminar synaptic strength following the MD 1d (left) and MD 4d (right), respectively. Data in panels are presented in mean ± s.e.m., the dots represent the connected synapses. See statistical data in Table S3.

Given that layer 2/3 PCs and PV neurons receive strong excitatory drive from layer 4^37^, we further examined the effect of MD on these inter-laminar inputs. We placed a stimulating electrode in layer 4 and measured evoked EPSCs in recorded PCs in layer 2/3, thereby generating the input-output curves with increasing stimulation intensities (**Figure 3C**). We found that only 4-day MD reduced the strength of these intra-laminar excitatory inputs onto PCs (*p* = 0.002, two-way ANOVA; **Figure 3D**), but not 1-day MD (*p* = 0.36; **Figure 3D**). Furthermore, the intra-laminar synaptic inputs onto layer 2/3 PV-INs were unchanged by either 1-day or 4-day MD (*p* = 0.28 and *p* = 0.10, respectively; **Figure 3E**).

Together with MD-effects on local excitatory synapses, these results systematically demonstrate an experience-dependent, laminar-specific reconfiguration of specific synaptic pathways in intracortical excitatory-inhibitory microcircuits over time. In general, 1-day MD preferentially modified excitatory transmission in layers 2/3 and 4, whereas 4-day MD selectively weakened local PV-PC inhibitory synapses and inter-laminar layer 4-2/3 excitatory transmission.

### 4-day MD consistently reconfigures disinhibitory microcircuits in both layers

We further examined the effects of MD on local inhibitory-to-inhibitory (I-I) or disinhibitory circuits among three recorded subtypes of INs with a connection probability of more than 10%. In layer 2/3, PV-INs not only form inhibitory connections to neighboring PV-INs, but also innervate SST-INs and VIP-INs (**Figures 1B** and **1C**). However, we found that neither 1-day nor 4-day MD significantly altered the strength (or uIPSCs) at these PV-PV (*p* = 0.86 and *p* = 0.26, respectively, Mann-Whitney *U* test), PV-SST (*p* = 0.30 and *p* = 0.87, respectively), and PV-VIP (*p* = 0.77 and *p* = 0.55, respectively) synapses (**Figures 4B**–**4D**). Given SST-INs preferentially target other IN subtypes^21^, we found that 1-day MD did not affect the strength of SST-PV (*p* = 0.16) and SST-VIP (*p* = 0.17) synapses, while 4-day MD enhanced the transmission at both SST-PV (*p* = 0.002) and SST-VIP (*p* = 0.009) synapses substantially (**Figures 4E** and **4F**). The latter synaptic strengthening was accompanied by increased connection probability at SST-PV synapses following 4-day MD (*p* = 0.018, Chi-square test; **Figure S3J**). In contrast, VIP-SST synaptic efficacy remained unaltered by MD at both time points (*p* = 0.096 and *p* = 0.69 for 1-day and 4-day MD, Unpaired *t* test and Mann-Whitney *U* test, respectively; **Figure 4G**). Thus, these results demonstrate that MD induces temporally delayed, synapse-specific plasticity in disinhibitory microcircuits of layer 2/3, with effects manifesting selectively at SST-PV and SST-VIP connections following prolonged (4-day) deprivation.

**Figure 4.**
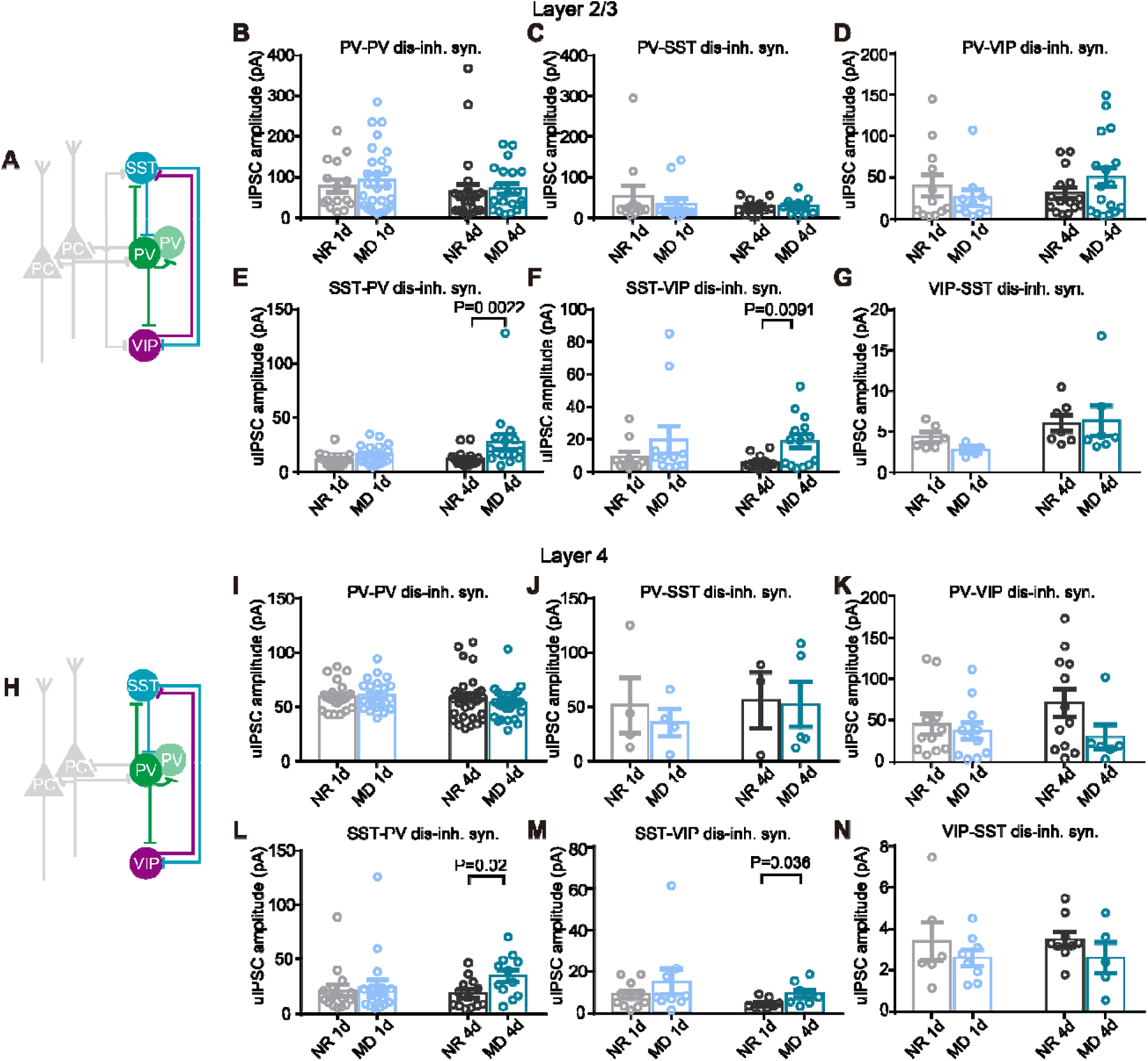
Identical reconfigurations of distinct disinhibitory synapses in both layers by 4-day MD. (A) A diagram illustrating the examined disinhibitory synapses (in colors) in the layer 2/3 microcircuit. (B-G) Changes of synaptic strength of various disinhibitory synapses among PV-INs, SST-IN and VIP-INs in the layer 2/3 microcircuit by MD 1d and MD 4d. The examined disinhibitory synapses included: PV-PV (B), PV-SST (C), PV-VIP (D), SST-PV (E), SST-VIP (F), and VIP-SST (G) disinhibitory synapses. (H) A diagram illustrating the examined disinhibitory synapses (in colors) in the layer 4 microcircuit. (I-N) Changes of synaptic strength of various reciprocal disinhibitory synapses among PV-INs, SST-INs and VIP-INs in the layer 4 microcircuit by MD 1d and MD 4d. The examined disinhibitory synapses included: PV-PV (I), PV-SST (J), PV-VIP (K), SST-PV (L), SST-VIP (M) and VIP-SST (N) disinhibitory synapses. Data in panels are presented in mean ± s.e.m., the dots represent the connected synapses. See statistical data in Table S4.

Interestingly, we observed the same effects of MD on modifying specific synapses in layer 4 disinhibitory microcircuits. Consistent with the above findings in layer 2/3, 4-day MD, but not 1-day MD, selectively potentiated transmission of SST-PV (*p* = 0.90 and *p* = 0.02 for 1 and 4-day MD, Mann-Whitney *U* test and Unpaired *t* test, respectively) and SST-VIP (*p* = 0.6 and *p* = 0.036, respectively) synapses (**Figures 4L** and **4M**). Conversely, all other disinhibitory synapses, including PV-PV, PV-SST, PV-VIP and VIP-SST synapses, were not affected by either 1 or 4-day MD (**Figures 4I**–**4N**)

Thus, these findings clearly show that prolonged (4-day) MD induces a conserved plasticity in disinhibitory microcircuits across cortical layers, specifically expressed as potentiation at SST-PV and SST-VIP synapses. Crucially, this suggests that the disinhibitory microcircuit is governed by a generalized, experience-dependent plasticity rule.

### MD-induced visual response plasticity of layer 2/3 PC, PV-, SST- and VIP-INs *in vivo*

Our above extensive *ex vivo* recordings have thus provided a comprehensive spatiotemporal map of experience-induced reconfigurations of visual cortical microcircuits during the CP. We next sought to determine how these systematic reconfigurations yield an *in vivo* plasticity of visual spiking activity in developing V1 neurons at early and late stages, which were reported previously^38,39^.

To address this question, we performed *in vivo* single-unit recording from PCs and PV-, SST-, VIP-INs in V1M layer 2/3 of anesthetized transgenic mice, under a two-photon laser scanning microscope (see Methods; **Figure 5A**). We measured the spiking rates of both the spontaneous activity, recorded during presentation of a blank screen, and the visually evoked activity responding to drifting gratings for each neuronal subtype (**Figure 5B**). Agree with a previous finding^40^, neither 1-day or 4-day MD modified the rates of visually evoked activity of excitatory PCs (**Figure 5C**, right). In contrast, 4-day MD, but not 1-day MD, significantly reduced the spontaneous spiking rate of these PCs (**Figure 5C**, left). Similar plasticity results were also observed in layer 4 PCs (**Figure S5B**), which accompanied with higher orientation selectivity **(Figure S6C**).

**Figure 5.**
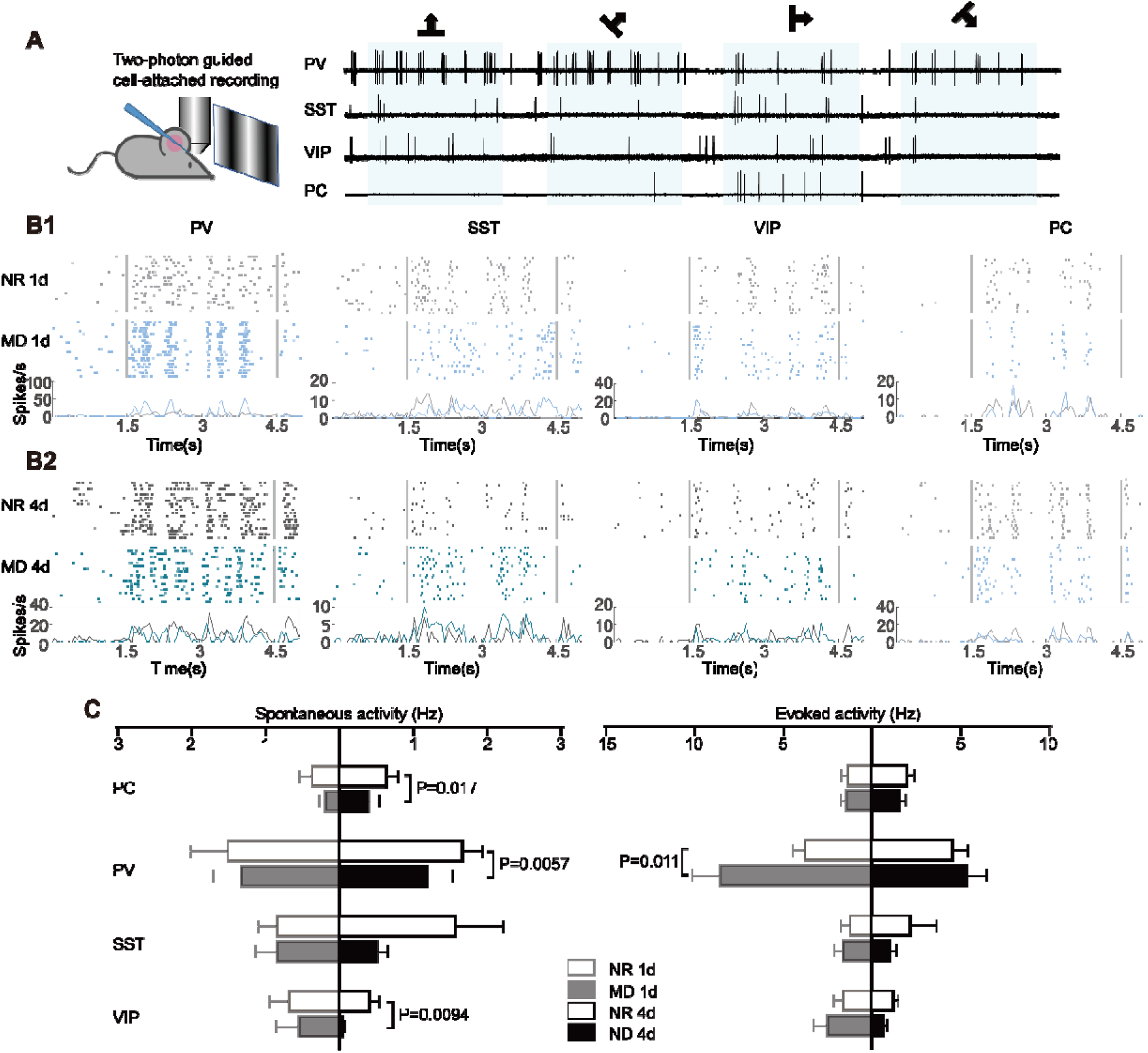
The effects of MD on spontaneous activity and evoked activity for PCs, PV-, SST-and VIP-INs in visual cortex layer 2/3. (A) Experimental setup showing two-photon guided recording for PV-, SST- and VIP-INs with the screen for visual stimulation (left). Visual response of various neurons in layer 2/3 for drift gratings with 4 orientations, the shaded areas indicate the period of presenting drifting grating (right). (B1) Representative visual response to preferential drifting gratings across 20 trials (top) and average visual response (bottom) of PCs, PV-, SST- and VIP-INs in the age-matched NR 1d (gray) and MD 1d (blue) mice, respectively; the bars represent spikes. (B2) Similar representative visual response (top) and average visual response (bottom) in the age-matched NR 4d (black) and MD 1d (cyan) mice, respectively. (C) Left: modulation of spontaneous activity by MD 1d and MD 4d for PCs, PV-, SST- and VIP-INs in the V1 layer 2/3. Right: modulation of evoked activity by MD 1d and MD 4d for PCs, PV-, SST- and VIP-INs in the V1 layer 2/3. Data in panels are presented in mean ± s.e.m., the dots represent the connected synapses. See statistical data in Table S5.

Furthermore, we found that 4-day MD did not alter visually evoked activity in any of the examined interneuron types (PV, SST, VIP). However, 1-day MD selectively and transiently increased the visually evoked response of PV-INs (**Figure 5C**, right). In contrast to evoked activity, 4-day MD significantly decreased the spontaneous activity rates of both PV- and VIP-INs, while 1-day MD did not affect the spontaneous firing of any INs (**Figure 5C**, left).

These *in vivo* recordings reveal neuronal subtype- and stage-specific plasticity of visual-related spiking activity in V1M neurons during MD. Notably, PV-INs uniquely exhibited transient potentiation of visually evoked responses after 1-day MD, while 4-day MD triggered a widespread reduction in spontaneous activity across all neuronal subtypes except SST-IN.

### A physiologically-based circuit model recapitulates cortical response plasticity

To bridge our observed specific synaptic microcircuit changes with the measured *in vivo* neuronal activity plasticity, we constructed a circuit model of V1M layer 2/3 (**Figure 6A**). The model incorporated the synaptic connection probabilities and strengths derived from our *ex vivo* measurements, and the PC was modelled with two compartments: a somatic compartment receiving bottom-up inputs (e.g. excitatory inputs from layer 4) and perisomatic inhibition from PV-INs as well as a dendritic compartment receiving top-down inputs and dendritic inhibition from SST-INs and other PCs. All three types of INs (PV, SST, VIP) were modelled as point neurons connected in a recurrent microcircuit (see Methods). This circuit network was implemented as a linear dynamical system designed to allow each neuron to converge to a stable firing rate over time (**Figure 6B**).

**Figure 6.**
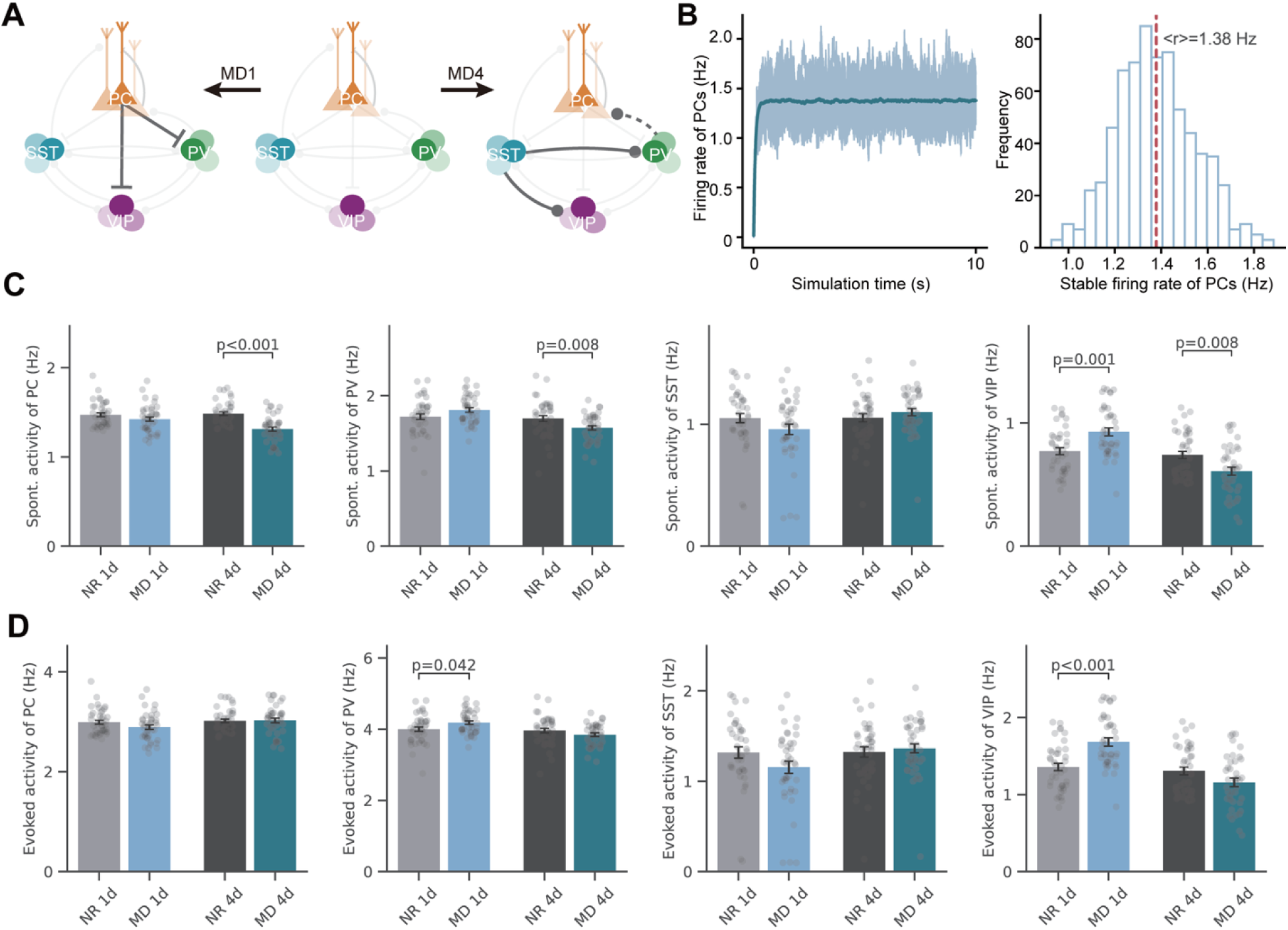
A microcircuit model accurately predicts neuronal activity changes. (A) The two-compartment microcircuit model with PCs (orange), PV (green)-, SST (blue)- and VIP (purple)-INs. Solid dark lines represent synapses with increased synaptic strength after MD and dash dark lines represent synapses with decreased synaptic strength after MD. (B) PCs’ activity as a function of simulation time (left). Light blue lines mark 20 different PCs randomly selected from the microcircuit and the dark blue line marks the mean activity of the PC population. The histogram of PCs’ activity after the network converged to a stable state (right). (C) Changes of spontaneous activity of PCs, PV-, SST- and VIP-INs with synaptic reconfiguration in the model. Mann-Whitney U tests were based on randomly sampling 3 cells from each of the 10 simulations, resulting in 30 cells in total. (D) Same as (C) but plotted under evoked condition.

We then quantitatively integrated our *ex vivo* findings of MD-induced synaptic locus- and stage-specific synaptic modifications in layer 2/3 microcircuits (1) enhanced PC-PV and PC-VIP excitatory transmission after 1-day MD, (2) weakened PV-PC inhibitory synapses and excitatory layer 4 inputs onto PCs after 4-day MD, and (3) enhanced SST-PV and SST-VIP disinhibitory synapses in layer 2/3 – to examine how these specific synaptic changes influence both spontaneous and visually-evoked activity from subtypes of neurons in layer 2/3. The model successfully recapitulated key *in vivo* results of spiking activity (**Figure 6C** and **6D**): 1-day MD selectively increased the evoked activity of PV-INs (*p* = 0.042, Mann-Whitney U test) whereas 4-day MD decreased the spontaneous activity of both PC (*p* < 0.001), PV-INs (*p* = 0.008) and VIP-INs (p=0.008). In parallel, the model revealed homeostatic circuit dynamics, as the spontaneous and evoked activities of many other neurons remained stable despite microcircuit reconfigurations. For example, the enhanced excitatory transmission onto PV- and VIP-INs during 1-day MD did not alter spontaneous network activity. This homeostasis arose might be due to increased interneuron activity suppressed PC firing, which in turn reduced excitatory drive back onto the interneurons, forming a stabilizing feedback loop.

### Modeling predicts a vital role of disinhibitory microcircuits in mediating plasticity

Furthermore, we utilized the microcircuit model to systematically dissect the contribution of individual synapses to the neuronal activity changes induced by MD. We quantified the impact of each type of synapses by updating one at a time and assessing how well the resulting model reproduced the activity of the full model (**Figure 7A**). This similarity was expressed as a reproduction index (RI), where a value of 1 indicated a perfect replication by only changing one type of synapse (see Methods).

**Figure 7.**
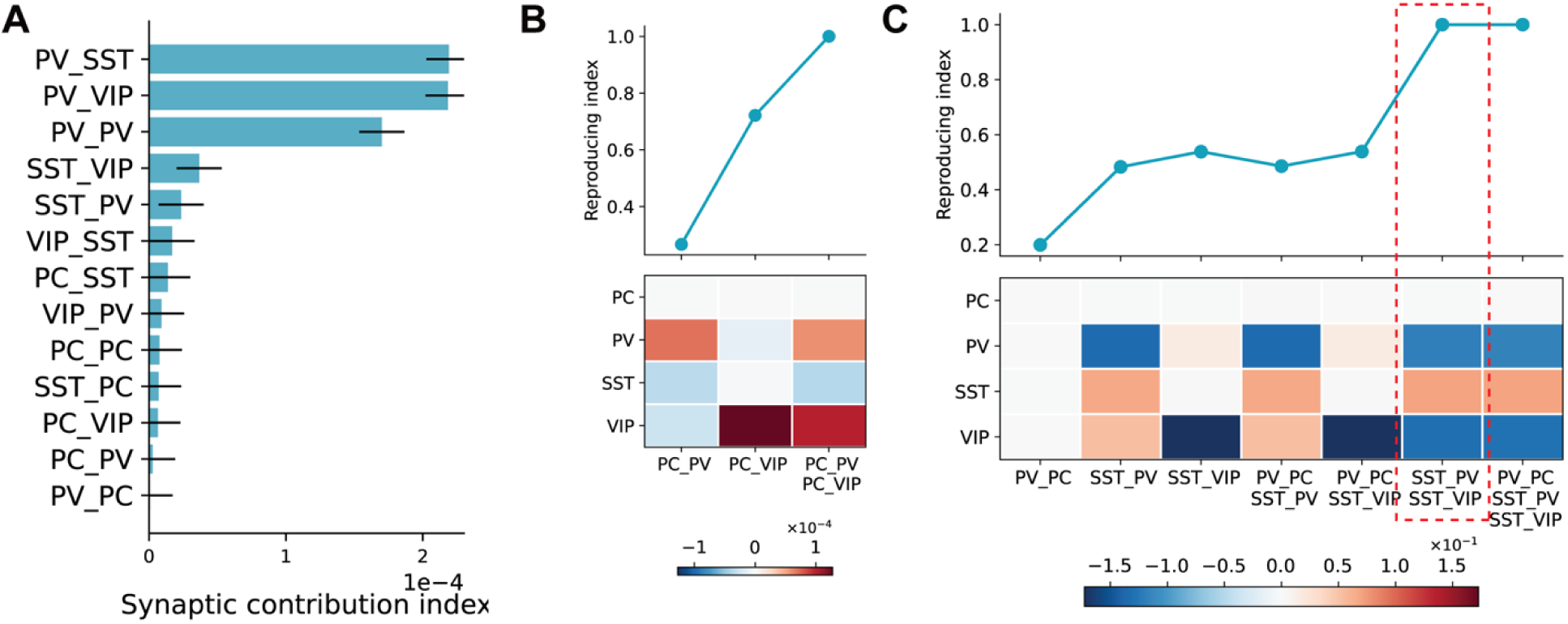
SST-PV and SST-VIP synapses cooperatively contribute to neuronal activity changes. (A) The synaptic contribution of 13 synapses in layer 2/3 microcircuit. (B) The replication index by changing a subset of the varied synapse in 1-day MD. (C) Same as (B) but for 4-day MD. Blue color marks the decrease of activity after reconfiguration of the synapses in the subsets, and red color marks the increase.

Our analysis revealed that for 1-day MD, changes of PC-VIP excitatory synapses (RI = 0.69) contributed more significantly to the observed activity change than that at PC-PV excitatory synapses (RI = 0.37). However, fully reproducing the 1-day MD-induced activity change required the cooperative action of both excitatory pathways (**Figure 7B**). In contrast, for 4-day MD, the two reconfigured disinhibitory pathways, SST-PV and SST-VIP synapses, were sufficient to almost entirely reproduce the full model’s activity change (RI = 0.97), underscoring the predominant role of disinhibition in the late expression of MD-induced activity plasticity (**Figure 7C**).

Beyond examining only those experience-reconfigured synapses, we extended our analysis to manipulate individual synapses in the model to gauge its general influence on the microcircuit functional activity. Intriguingly, the synapses that were modified by MD fell in the middle of the sensitivity spectrum (**Figure 7A**), indicating that they were neither highly sensitive nor insensitive to activity changes. In contrast, the most and least sensitive synapses—those with the largest synaptic contribution indices (see Methods) for influencing activity—remained unchanged.

In summary, while prior research has preferentially focused on isolated synaptic motifs, our modeling results indicate that synaptic influences on cortical functional activity in a cooperative manner. More specifically, we have identified the SST-PV and SST-VIP disinhibitory pathways as major drivers for neuronal activity changes or plasticity in layer 2/3 microcircuits following sensory deprivation.

## Discussion

Our study has defined a sequential and cell-type-specific microcircuit mechanism underlying experience-critical period plasticity of developing V1M. We demonstrate that brief MD (1-day MD) transiently reconfigures excitatory synapses onto inhibitory PV-INs and VIP-INs selectively in cortical layers 2/3. In contrast, prolonged deprivation (4-day MD) primarily strengthens inhibitory transmission from SST-INs onto PV-INs and VIP-INs in both layers, highlighting a critical role of disinhibitory microcircuits in the experience-dependent plasticity. Critically, modelling computations, based on our empirical data, suggest this coordinated sequence of synaptic changes—first excitatory, then disinhibitory circuits—is necessary to mediate the differential, experience-dependent plasticity of visual responses in layer 2/3 V1M neurons *in vivo*. Together, these findings provide a unifying microcircuit framework for elucidating how distinct subnetworks are sequentially recruited to orchestrate cortical plasticity during the CP.

### Conserved motifs across layers

Many studies systemically described connectivity between subtypes of neurons in visual cortex by electrophysiological recordings or purely electron microscopy reconstruction methods. As for the disinhibitory synaptic network, we confirmed the similar motifs prominently in intracortical microcircuits, that PV-INs tend to inhibit each other rather than inhibit neighboring INs, and SST-INs broadly inhibit other subtypes of INs except themselves, whereas VIP-INs preferentially innervate SST-INs^25,27,28^. Moreover, we notice additional disinhibitory motifs, including PV–VIP, SST–VIP, and PV–SST connections, which have received limited attention in previous study^18,41,42^, and the recurrent connections among PV–SST and VIP–SST INs are even more prevalent than those among PCs in the cortex (**Figure 1**), these reciprocal connections among subtypes of INs we observed may have important implications for cortical function and plasticity^43^.

Although the spatial distribution of INs varies strongly by layers, our study suggests that the disinhibitory microcircuits are conserved across layers. This finding conflicts with the previous study that SST-INs predominantly innervate neighboring PCs in layer 2/3, while they primarily target fast-spiking PV-INs in layer 4^24^. This discrepancy may arise from use of different transgenic mouse lines (*X94* vs. *SST-cre*) or different cortex regions (somatosensory cortex vs. visual cortex). Alternatively, it suggests that broad molecular classifications (e.g., PV, SST, VIP and others) are insufficient to capture the full heterogeneity of cortical INs. Indeed, recent studies on spatial transcriptomic analysis reveal the laminar organization of eight SST-INs subtypes, and subtypes of SST-INs form cell-type-specific cortical circuits^44^. Future work could leverage single cell RNA-seq data to generate intersectional transgenic tools for labeling specific subpopulations of excitatory and inhibitory neurons, enabling a more refined dissection of laminar circuit connectivity^45,46^. Moreover, the emergence of new technologies, like multiphoton optogenetic stimulation, offers new opportunities to dissect laminar circuit connectivity in cortex^47^.

### Temporal reconfigurations of microcircuits by experience in developing V1M

Numerous studies have examined the effects of MD on reconfiguring synaptic microcircuits of developing V1, largely utilizing *ex vivo* measurements. Collectively, this body of work indicates that the reconfiguration of intracortical synapses is a dynamic process with distinct temporal profiles, exhibiting layer- and synaptic-specific plasticity^8,12,16,19^. For example, prolonged MD (>3 days) has been shown to significantly enhance inhibitory synaptic transmission onto layer 2/3 PCs, whereas 1-day MD does not^16^. Other studies have reported that 2 and 6 days of MD induce opposite changes in excitatory inputs onto layer 2/3 PCs^39^. In addition to synaptic reconfiguration, visual deprivation drives progressive structural changes on presynaptic boutons and postsynaptic dendritic spines over days^48,49^. A recent study found that the developmental increase in bouton formation was disrupted after 3 days of MD, the postsynaptic spines remodeled quickly in response to MD in layer 2/3 PCs. The studies have shown the effect of temporal visual deprivation on structure and synaptic strengthens of intracortical connections^50^. The systematically and temporally resolved circuit mechanisms underlying experience-dependent plasticity in V1M have remained largely unclear. Focus on synaptic reconfigurations in layer 2/3 and 4 microcircuits by experience with more systematic and thorough approaches.

Here, we demonstrate (1) 1-day MD rapidly remodels specific excitatory synapses (**Figure 2**), and (2) 4-day MD drives changes in the same selective disinhibitory synapses across layers, suggesting a conserved circuit motif (**Figure 4**). Our studies establish a temporal framework for experience-dependent synaptic remodeling in V1M. Meanwhile, we systematically dissect the effects of MD on broad synapses and examine across layer 2/3 and layer 4 microcircuits and investigate how these circuit changes influence visual responsiveness at the systems level. A key question arising from the observation of MD-induced temporal synaptic reconfigurations, is how early-stage (1-day MD) synaptic modifications enable later-stage (4-day MD) plasticity. We propose a two-stage model in which experience-induced changes in intracortical excitatory synapses encode the initial sensory input during early MD, then early excitatory changes may gate, or constrain subsequent inhibitory and disinhibitory plasticity. It will be important to test whether computational circuit models can reproduce these sequential plasticity dynamics. Such modeling insights will be crucial for building a comprehensive framework for experience-dependent plasticity.

Our results show that 1-day and 4-day MD during the CP bi-phasically regulate the visually evoked spiking activity of layer 2/3 PV-INs in V1M (**Figure 5C**), suggesting the involvement of homeostatic plasticity. However, our observation of 1-day MD-induced increased activity is inconsistent with a previous report^39^, and this inconsistency may be attributable to difference in experimental condition, such as brain states (awake vs. anesthetized), visual stimuli (simple drifting gratings vs. natural scenes) and species (rat vs. mouse).

By comparing the synaptic reconfigurations between layer 2/3 and 4, we demonstrate that 1-day MD regulates the strength of PC–PV excitatory synapses in layers 2/3 and 4 in opposite directions (**Figures 2C** and **2H**). This layer-specific differential modification indicates that synaptic changes are not uniformly expressed across cortical layers. One explanation is rooted in the canonical thalamo-cortical pathway, in which thalamic afferents primarily drive layer 4 PCs before relaying activity to layer 2/3. These anatomical constraints create distinct input–output transformations in the two layers, potentially enabling parallel and independently regulated plasticity mechanisms. Layer 4 PV-INs circuits may be more directly influenced by changes in thalamic drive immediately after deprivation, whereas layer 2/3 PV-INs circuits may integrate both altered feedforward input and intracortical activity patterns. More broadly, these findings suggest that early experience-dependent plasticity of excitatory microcircuits is also expressed into layer-specific manner, which supports the previous studies^4,20,31^.

### Experience-dependent visual function plasticity in developing V1M and V1B

V1M in the mammalian V1 primarily processes visual inputs from the contralateral eye, and unlike the V1 binocular area (V1B), where ipsilateral eye input pathway is intermingled with the contralateral synaptic pathway^4^. Since homosynaptic LTD in the deprived-eye pathway, induced by loss of deprived-eye responsiveness, is present in both V1M and V1B following MD, multiple studies have reported similar synaptic reorganization in the two regions^12,13,39,40,51^. However, the absence of pronounced inter-ocular competition means that V1M lacks homeostatic plasticity in the non-deprived eye pathway like V1B^52^ ined in V1M should not be readily generalized to V1B. Moreover, our *in vivo* data demonstrate that PV-INs exhibit increased activity in the 1-day MD, conflicts with the research in V1B, which show decreased PV-INs activity evoked by both contra- and ipsi-lateral eye inputs^17^. Thus, validating these findings within the well-characterized V1B microcircuit is essential, particularly given the unresolved question of whether V1M and V1B have evolved distinct or parallel cellular mechanisms of plasticity.

### Disinhibitory microcircuits underlying the regulation of cortical plasticity

In the present study, we systematically assess the effects of MD on broad disinhibitory synapses^4^, which are mechanistically distinct from the commonly studied PC–IN microcircuits. Remarkably, experience-induced plasticity in SST-PV and SST-VIP disinhibitory synapses emerges as a conserved feature across both layers (**Figure 4**), suggesting that disinhibition may serve as a shared circuit module for regulating visually deprivation-induced changes. This cross-layer consistency implies that SST interneurons may occupy a central position in orchestrating experience-dependent shifts in excitation–inhibition balance by selectively modulating inhibitory subnetworks. Such a motif could provide a flexible means of shaping plastic windows while preserving overall network stability. Future work will be needed to determine whether these disinhibitory plasticity mechanisms extend to deeper layers of V1 or to other sensory cortices, which would support the broader idea that disinhibitory microcircuits act as a canonical scaffold for experience-dependent circuit reorganization. In order to elucidate how distinct synapses interact to shape neuronal activity in layer 2/3, we employed an empirical data-based circuit model and well predicted plasticity of neuronal spiking activity induced by the brief or prolonged MD. We noted that the model’s prediction for VIP-INs activity (**Figure 6**) did not fully align with the experimental observations (**Figure 5**). This discrepancy arises partly from the fact that our simplified microcircuit model omitted several important long-range inputs to layer 2/3, such as thalamic, cortico-cortical, and top-down neuromodulatory projections^18,53–55^. The model’s overall predictions are consistent with our *in vivo* measurements, thereby providing a mechanistic link between observed systematic synaptic plasticity and neuronal functional neuronal plasticity. Intriguingly, our simulations defined that specific disinhibitory synapses from SST-INs onto PV-INs and VIP-INs play an active and dynamic role in plasticity (**Figure 7**).

This agrees with previous studies in which disinhibition was shown to regulate the CP plasticity in developing V1B^17,18,56^. A recent study demonstrated that cholinergic inputs selectively recruit SST-INs and subsequently modulate their activity specifically during the CP, a process that then enhances dendritic inhibition in PCs via activation of SST-PC inhibitory synapses along with concurrently disinhibiting somatic compartments through the SST-PV-PC pathway^18^. These findings collectively support a model in which disinhibition enables a transient ’window of opportunity’ for experience-dependent plasticity.

Furthermore, our model also predicted that the SST-PV and SST-VIP disinhibitory synaptic motifs operate cooperatively to govern neuronal activity changes. This represents a significant departure from the prevailing view that a given synaptic motif implements a defined plasticity rule and function^17,18,57^. While several reviews have proposed that a more complex microcircuit may underlie critical period plasticity^4^, our findings demonstrate that the underlying circuit mechanisms are indeed more intricate than previously characterized. Specifically, we show that this plasticity requires the cooperative action of disinhibitory motifs formed by the three major inhibitory neuron types. A further exploration of these disinhibitory mechanisms in V1 may yield novel insights into experience-dependent plasticity and could potentially inform strategies to modulate plasticity in adult brain.

## Resource availability

### Lead contact

Requests for further information and resources should be directed to and will be fulfilled by the lead contact, Xiaohui Zhang.

### Code availability

Code for reproducing results from the microcircuit model is available at: https://github.com/ZilongJi/SynReconfigNet

## Acknowledgements

This work was supported by grants from the National Natural Science Foundation of China (32130043) and the Scientific & Technological Innovation (STI) 2030-Major Project (2022ZD0204900) to X-h. Z., and by the National Natural Science Foundation of China (32400870) and the Fundamental Research Funds for the Central Universities (2243300002) to Z.-w.S.

## Author contributions

X-h.Z., L.Y. and Z-w.S. conceived and designed the experimental procedures. M.H. provided trans genetical mice. L.Y. and M-z.M. conducted the slice recording experiments. L.Y. and Z-w.S. conducted immunostaining experiments. Z-w.S. and F-c.W. conducted in vivo recording experiments. Z-l. J. and S.W. performed modeling computation. L.Y. and Z-w.S. conducted all the data analysis. X-h.Z., L.Y., Z-w.S. and Z-l.J. wrote the manuscript with the comments from all other authors.

## Supplemental information

Document S1. Figures S1–S6, Table S1

Document S2. Tables S2–S11

## Methods

### Key resources table

| REAGENT or RESOURCE | SOURCE | IDENTIFIER |
| --- | --- | --- |
| **Antibodies** |  |  |
| fluorophore-conjugated streptavidin | Invitrogen | S32354;<br>RRID:AB_2315383 |
| DAPI | Beyotime | C1002;<br>RRID:AB_3675433 |
| **Experimental models: Organisms/strains** |  |  |
| Mouse: wild type *C57BL/6J* | Charles River |  |
| Mouse: *Pvalb-IRES-Cre* | The Jackson Laboratory | Jax No. 008069;<br>RRID:IMSR_JAX:008069 |
| Mouse: *Sst-IRES-Cre* | The Jackson Laboratory | Jax No. 013044;<br>RRID:IMSR_JAX:013044 |
| Mouse: *Vip-IRES-Cre* | The Jackson Laboratory | Jax No. 010908;<br>RRID:IMSR_JAX:010908 |
| Mouse: Ai9, *ROSA26-CAG-tdTomato* | The Jackson Laboratory | Jax No. 007909;<br>RRID:IMSR_JAX:007909 |
| Mouse: B13, *PV-EGFP* | provided by Dr. J.Z. Huang |  |
| Mouse: GIN, *FVB-Tg (Gad-GFP) 45704Swn/J* | provided by Dr. J.Z. Huang |  |
| **Software and algorithms** |  |  |
| MATLAB R2020b | MathWorks | https://www.mathworks.com |
| pClampex (version 10.3) | Molecular Devices | https://www.moleculardevices.com/products/axon-patch-clamp-system |
| ImageJ (FIJI) | NIH | https://imagej.net/software/fiji/ |
| BrainPy | Wang et al.<sup>58</sup> | https://github.com/PKU-NIP-Lab/BrainPy |
| Python | Python Software Foundation | https://www.python.org |
| GraphPad Prism 10 | Graphpad Software | https://www.graphpad.com/scientificsoftware/prism/ |

## Experimental model and study participant details

Wild type C57BL/6J and six different lines of transgenic mice of postnatal days 27-33 (P27-33), either male or female, were used. The transgenic lines B13 (*PV-EGFP*) and GIN (*FVB-Tg (Gad-GFP*) 45704Swn/J) mice were generously provided by Dr. J.Z. Huang (CSHL), while the knock-in lines *Pvalb-IRES-Cre* (by S Arbor at FMI, Jax No. 008069), *Sst-IRES-Cre*, *Vip-IRES-Cre* (both by J. Z. Huang at CSHL; Jax No. 013044 and 010908, respectively) were crossed with the tdTomato-reporter line Ai9 (*ROSA26-CAG-tdTomato*, by H.K. Zeng at Allen Brain Institute, Jax No. 007909) to selectively label the parvalbumin (PV)-, somatostatin (SST)- and vasoactive intestinal polypeptide (VIP)-expressing inhibitory interneurons (INs), respectively. To achieve simultaneous labeling of either two subtypes interneurons among the above three IN subtypes, *Pvalb-IRES-Cre::Ai9* or *Vip-IRES-Cre::Ai9* mice were further crossed with the *GIN* mice, whereas *Vip-IRES-Cre::Ai9* were crossed with the *B13* mice. All mice were housed in the animal house facility on a 12 hr light/dark cycle and the protocols of all animal experiments were approved by the State Key Laboratory of Cognitive Neuroscience & Learning at Beijing Normal University (IACUC-BNUNKLCNL-2020-04).

## Method details

### Monocular vision deprivation

The eyelid suture was conducted to an eye in mice at P27 or P28, under isoflurane anesthesia (0.5–2% in oxygen) and body temperature maintenance at 37 (RWD Life Science, China) to achieve monocular deprivation (MD) of visual inputs, following a method described previously^5^9. After the recovery from anesthesia, mice were exposed to normal 12 hr light/dark cycles in their home cages. The MD treatment lasted for either 1 day or 4 days, during which animals were daily checked to ensure the sutured eye closed and uninfected. After the MD, the sutured eye was reopened right before the *in vivo* single-unit recording or the brain slice preparation.

### Preparation of visual cortical slices

The detailed procedure of preparing mouse cortical slices followed a method described in our previous studies^34,60^. In brief, mice were anesthetized by intraperitoneal injection of sodium pentobarbital (50mg/kg of body weight). The brain was dissected rapidly and placed into the ice-cold, oxygenated (95% O_2_ and 5% CO2) artificial cerebral spinal fluid (aCSF), containing (in mM) 125 NaCl, 1.25 NaH_2_PO_4_, 2 CaCl_2_, 3 KCl, 2 MgSO_4_, 1.3 Na^+^-ascorbate, 0.6 Na^+^-pyruvate, 26 NaHCO_3_, 11 dextrose (pH 7.25, 300 mOsm). Coronal cortical slices at 300 µm thickness, containing the primary visual cortex (V1) contralateral to the sutured eye, were prepared using a Leica vibratome (VT-1200S, Leica, Germany) and transferred into a custom-made incubation beaker filled with oxygenated aCSF at 34 for 30min, followed by an incubation at room temperature (20–22 ) for more than 30min before recording.

### Whole-cell recording in cortical slices

Cortical slices were placed in the recording chamber with a perfusion of the oxygenated aCSF at a rate of 2 ml/min and the solution temperature was maintained at 31 ± 1 (TC-344B, Warner Instrument Corporation, USA) during the whole-recording procedure. The monocular zone (V1M) in the V1 slices was identified with anatomical characteristics of relatively dense neuronal processes and the white matter under a 5x objective lens of an upright microscope (FN1, Nikon, Japan), equipped with the differential interference contrast (DIC) optics and infrared Rolera-XR CCD camera (QImaging, USA). Simultaneous whole-cell recordings were made from pairs of excitatory pyramidal cells (PCs) or inhibitory PV-, SST-, VIP-INs within the layer 2/3 or the layer 4 in the V1M, with an Axon Multi-Clamp 700B amplifier (Molecular Devices, USA). Distances between the two recorded neurons were normally within 100 µm. The recording borosilicate micropipettes (BF 120-69-15, Sutter Instrument, USA) were made by a Sutter P-1000 puller and filled with either a standard internal solution containing (in mM) 130 K^+^-gluconate, 20 KCl, 10 HEPES, 0.2 EGTA, 4 Mg_2_ATP, 0.3 Na_2_GTP, and 10 Na_2_-phosphocreatine (pH 7.25-7.35; 305 ± 5 mOsm) for recording unitary excitatory postsynaptic currents (uEPSCs), or a 60-mM high chlorine internal solution (by increasing KCl in replace of K-gluconate) for recording unitary inhibitory postsynaptic currents (IPSCs). The resistances of micropipettes ranged 3-6 MΩ. In some experiments, biocytin (0.2–0.3%, Sigma) was added to the internal solution for post-hoc immunostaining reconstruction of the morphology of recorded neurons.

To assay the efficacy and dynamics of unitary synaptic transmission of various intralaminar excitatory or inhibitory synapses, a train of 5 current pulses at 20 Hz (1ms duration, 1.2–3 nA) was delivered intracellularly into the presynaptic neuron to faithfully elicit action potentials under the current-clamp mode, while evoked uEPSCs or uIPSCs were recorded in the postsynaptic neuron under the voltage-clamp mode and averaged from 30 sweeps. For the trans-laminar excitatory synapses from layer 4 to layer 2/3, a field stimulation of 5 pulses (1-ms duration) at 20 Hz was made with a concentric tungsten electrode (Biospikes, China), placed approximately 60 µm below the slice surface within layer 4, and evoked EPSCs in the layer 2/3 PCs or PV-INs, located 50–150µm vertically from the stimulation site, were recorded under the voltage-clamp mode in the presence of GABA_A_ receptor antagonist picrotoxin (PTX, 50 µM) in the bath solution. The field-stimulation intensities were increased at 6 steps from 10 µA to 100 µA, and each intensity was repeated for 8 sweeps. Electrical signals were low-pass filtered at 10 kHz (for action potentials) or at 4 kHz (for EPSCs or IPSCs), digitized at 20 kHz by the Digidata 1440A (Molecular Devices, USA) and acquired by the pClampex (version 10.3) in a computer. Data recorded from neurons showing access resistance (Ra) > 30 MΩ, and resting membrane potential > -60 mV were excluded for further analysis.

### Visual stimulation

Visual stimuli were generated by a custom-made software written using LabVIEW (National Instruments, USA) and MATLAB (Mathworks, USA) as described by our previous study ^59^. The stimuli were presented to the mice using a cathode ray tube (CRT) monitor (G520, SONY, Japan), placed 20 cm in front of the sutured eye. The receptive fields of individual recorded cells were roughly measured by flashing bright squares in an 8×8 location grid on the screen. Individual squares (9° x 11°, 100% contrast) in the grid were presented in a pseudo-random sequence at 20 Hz with 50 repetitions. The full-field drifting gratings (100% contrast) with 8 different orientations, at 0.02 cpd (cycle per degree) spatial frequency and 2 Hz temporal frequency, were used to measure the preferred orientation of recorded V1M neurons. To determine visually driven activity, full-screen drifting gratings with optimal orientation were presented with 20 repetitions. Each trial consisted of 3 s drifting gratings and 7-8 s gray screen (mean luminance). Spontaneous activity was measured by 1 min black screen.

### Single-unit recordings *in vivo*

Extracellular single-unit recording of V1 neuron spikes evoked by visual stimuli in anesthetized wild-type mice followed our previous studies^59,60^, with some modifications for the V1M recording. In brief, mice under the anesthetized state by ketamine (50 mg/kg) and medetomidine (0.6 mg/kg, i.p. injection) were mounted on a custom-made stereotaxic device and their body temperature was maintained at 37 with a controlled temperature pad (RWD Life Science, Chinese). A 1 mm ×1 mm craniotomy was made above the V1M region contralateral to the previously sutured eye, plus a black eye-shield covering the non-sutured eye during the whole recording procedure. The borosilicate glass micropipettes, filled with the normal saline containing 0.2–0.3% biocytin (w/v, with 4–8 MΩ resistance), were used to record single-unit spike from V1M neurons, which resided in the layer 2/3 and layer 4 at depths of 150–350 µm and 400–600 µm under the pia (Chen et al., 2014), respectively. The membrane potential signals were amplified by an Axon Multi-Clamp 700B amplifier (Molecular Devices, USA), low pass filtered at 5 kHz, digitized at 10kHz with an Axon Digidata 1440A convert board (Molecular Devices, USA), and then acquired by the pClampex (version 10.3) to a computer for further analysis by the custom program in the MATLAB (Mathworks, USA).

### Two-photon guided cell-attached recording *in vivo*

The two-photon laser imaging-guided cell-attached recording was conducted to measure spontaneous and evoked spiking activity specifically from PV-, SST- and VIP-INs in the *Pvalb-IRES-Cre::Ai9*, *Sst-IRES-Cre::Ai9* and *Vip-IRES-Cre::Ai9* mice, respectively, following a method established in our previous study^60^, also with some modification for V1M recording. Similarly, under the ketamine/medetomidine anesthesia, a 1 mm ×1 mm craniotomy was made above the V1M region contralateral to the previously sutured eye. After the dura was moved, dental was used to surround the craniotomy to form a recording chamber which was filled with the Ringer’s solution containing (in mM) 116 NaCl, 2.9 KCl, CaCl_2_ 1.8, HEPES 5 (pH 7.2–7.4).

Glass micropipettes were filled with the normal aCSF containing 50 µM Alexa Fluor 488 (pH 7.3, 300 mOsm), with micropipette tip resistance of 4– 10 MΩ, to visualize the micropipette advancing in the brain during the two-photon laser imaging *in vivo*.

Cell-attached recordings were targeted to a specific subtype of INs within the V1M layer 2/3, labeled by tdTomato expression in three different transgenic mice, under an Olympus two-photon laser-scanning microscope (FV1200MPE, Olympus, Japan) equipped with an ultrafast Ti: Sapphire laser (Mai Tai-DeepSee, Spectra Physics, USA). The 960-nm laser beam was set to excite tdTomato inside the INs and Alexa 488 (50μM, Molecular Probes) inside the micropipette. An XL-PLAN 25x water-immersion objective (N.A. 1.05, Olympus, Japan) was used to continuously image cortical neurons and the micropipette when the micropipette was advancing to targeted INs, during which a positive pressure of 0.2 psi was applied to the micropipette until it touched the cell membrane. The latter step was indicated by a dramatic drop in the magnitudes of current responding to the test voltage pulse (5 mV, 100 Hz). Immediately, the positive pressure was released and then a small amount of suction was applied to form a loose seal of resistances ranging from 20 MΩ to 100 MΩ. Such loose-seal contact of the micropipette with the target cell membrane allowed to record spikes from a single IN without rupturing the neuronal membrane, using an Axon Multi-Clamp 700B amplifier (Molecular Devices, USA). Spike signals were low-pass filtered at 5 kHz, digitized at 10 kHz with an Axon Digidata 1440A convert board (Molecular Devices, USA), and acquired by the pClampex (version 10.3) in a computer for further analysis by a custom program in MATLAB (Mathworks, USA). During the surgery and recording, the eye cream was applied to prevent the eyes from drying.

### Neuronal morphology reconstruction and confocal imaging

To reconstruct the neuronal morphology and further determine the laminar location of *in vivo* recorded neurons, biocytin was infused into the neuron via the internal solution during the recording. After the recording, the mouse was fixed by transcardial perfusion with 4% paraformaldehyde (PFA, in 0.01 M phosphate buffer saline PBS). The fixed brain was dissected and further fixed with PFA at 4 overnight, followed by being coronally cut into 70 µm sections with a vibratome. Cortical sections were then treated with 0.5% Triton X-100 and 10% bovine serum albumin (BSA) in PBS for 1 hour. After being washed by PBS for 3 times, they were incubated with fluorophore-conjugated streptavidin (1:1000, Invitrogen) for 2 hours at room temperature. Meanwhile, cell nucleus staining by DAPI (1:2500, Beyotime) was made to visualize the laminar structure of the V1 based on differential densities of stained cortical cells. Fluorescence images of cortical tissue and biocytin-infused neurons were acquired by a confocal microscope (A1, Nikon, Japan) with a Plan Apo 20× objective (N.A. 0.75, Nikon, Japan). Neuronal morphology was further reconstructed manually with Image J (National Institutes of Health, USA).

### The microcircuit model

We built a rate based two-compartment model to understand the detailed dynamics under MD, which consists of an excitatory population PCs, and three inhibitory populations corresponding to PV, SST and VIP interneurons. The firing activity of a PC with index *i* evolves according to the following dynamics:

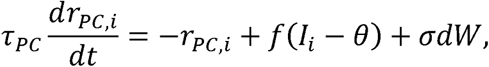

where *τ_PC_* is the time constant of the firing rate of PC, *I_i_* is the total somatic input and *θ* is the rheobase which resembles the smallest injected step current that results in one action potential. *f*(*x*) is the activation function:

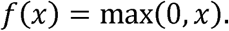

*σ* is the noise strength and *dW* is a Gaussian white noise with unit variance applied to the PC.

The total synaptic input *I_i_* is determined by both the somatic and dendritic synaptic events, as well as the potential dendritic calcium spiking events, which is written as:

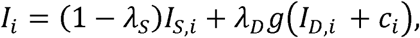

with *λ_S_* and *λ_D_* representing the proportion of currents leaking away from the soma and dendrites, respectively. *I_S,i_* and *I_D,i_* are the somatic and dendritic synaptic events, respectively. *c_i_* represent the calcium dendritic event in the PC neuron. *g*(*x*) is a nonlinear function describing the integration effect in the dendrites which is expressed as *g*(*x*) = max (0, *x*). The three events are described in detail as follows:

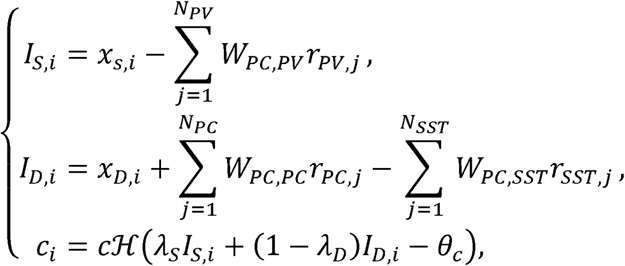

where *x_S,i_* and *x_D,i_* denote the external bottom-up input to the soma and top-down input to the dendrite, respectively. *r_PV,j_* represents the firing rate of the *j_th_* PV neuron, which mainly inhibits the peri-somatic regions and the basal dendrites of PCs. *r_SST,j_* represents the firing rate of the *j_th_* SST neuron which mainly inhibits the distal dendrites of PCs. *r_PC,j_* represents the firing rate of the *j_th_* PC which excites the distal dendrites of PCs. *W_PC,PC_*, is the recurrent excitatory strength, while *W_PC,PV_*, and *W_PC,SST_*, are the synaptic connection strength from the PV neurons and SST neurons to the PCs, respectively. *N_PC_*, *N_PV_* and *N_SST_* represent the number of PC, PV and SST neurons which have connections with the *i_th_* PC. *H*(*_x_*) is the Heaviside step function which gives 1 if *x* > 0 and 0 if *x* ≤ 0. It models the emission of a calcium spiking event if the input amount gathering at the dendritic site (i.e., the summation of the synaptic input leaked from the soma *λ_S_I_S,i_* and the synaptic input remaining at the dendrites (1 − *λ_D_*)*I_D,i_*, exceeds a threshold *θ_c_*. The spike event is scaled by a gain factor *c* to model the amount of current produced by the calcium spike.

For the firing activity of an interneuron with index *i*, it evolves according to the following dynamics:

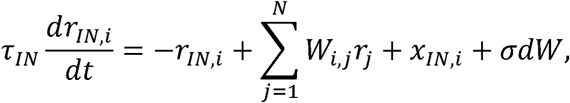

where *τ_IN_* is chosen to resemble *GABA_A_* time constant for all the interneurons. *x_IN,i_* is the external input drive. *W_i,j_* is the synaptic connection strength from other neurons, including PC, PV, SST and VIP. *σ* is the noise strength and *dW* is a Gaussian white noise with unit variance applied to the current interneuron.

To simulate the two-compartment model, we set the parameters based on our own data, as well as previous studies^61,62^. Both the probabilities and strengths of synaptic connections are adopted based on the data from our *in vitro* dual-patch whole cell recording experiments. Noteworthy, the original strength values are the postsynaptic currents (pA) recorded under the voltage-clamp configuration. To avoid the numerical instability during simulations, we scale down all the connection strengths by normalizing them with the largest value (i.e., 79 pA from PV to PC).

The time constants and *τ_PC_* and *τ_IN_* are set to 10 ms. The rheobase *θ* of pyramidal cells is set to 14 and the threshold *θ_c_*for generating dendritic spiking events is set to 28. *λ_S_*=0.31, *λ_D_*=0.27, and *c*=7. The external inputs to the network are set in order to match the mean firing rate of the four cell types measured in wild-type mice, which are different under the spontaneous and evoked conditions. Specifically, under the spontaneous condition, all cells within the same cell type share the same amount of input, which are *x_S,i_* =18.0, *x_D,i_* =5.0;, *x_PV,i_*=3.1, *x_SST,i_*=1.9, *x_VIP,i_* =1.4. Under the evoked condition, input values are *x_S,i_* =22.8, *x_D,i_* =10.0; *x_PV,i_* =7.1, , *x_SST,i_*=3.3, *x_VIP,i_* =2.8. Furthermore, for the 4d MD, we also observed that there is a decrease in the feedforward input from layer 4 to layer 2/3 in empirical data. Therefore, we decrease the bottom input *x_S,i_*, to 17.4 in the spontaneous condition, and to 20.7 in the evoked condition.

Simulation related parameters are set as follows: the number of PCs is 700, the number of PV-, SST- and VIP-INs are set to 100. The initial firing rate of all neurons are set to 0 at the beginning of the simulation, i.e., *r_PC,i_*=0 and *r_IN,i_*=0. The duration of simulation is set to 1000 ms, which is sufficient for the network to converge to the steady state. The integrating time step is set to 0.1 ms with a four-order Runge-Kutta method. All the simulations are carried out based on the BrainPy platform (https://github.com/PKU-NIP-Lab/BrainPy). For the summary of all the parameters described above, see Table S11.

## Quantification and statistical analysis

### Spontaneous and evoked spiking activity

For individual recorded neurons, the mean rate of its spontaneous or baseline spiking activity was measured by averaging spikes that occurred during the 1 min period of black screen. The evoked activity was calculated by averaging evoked spikes over the 3-sec drifting gratings with the preferential orientation after a subtraction from the mean baseline rate, thus it was presented as spikes per sec.

### Orientation selectivity

The orientation selectivity was quantified by a global measurement of orientation selectivity:

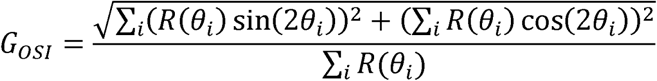

where *θ_i_* is the angle of the moving direction of the grating, and R(*θ_i_*) is the is the spike response (subtracted the baseline) at angle *θ_i_*.

### Statistics

sAll statistical analyses were performed using Prism (GraphPad Software) and The MATLAB R2020b (MathWorks). The unpaired t-tests were used to measure the significance of the difference between data sets if they followed a normal distribution (assayed by the Shapiro–Wilk test). Otherwise, non-parametric tests, the two-sided Mann-Whitney U test or the two-way analysis of variance (ANOVA) were used for statistical comparisons between independent groups. The chi-square test was conducted to determine the significance of the connectivity probability difference between groups. All data are presented as mean ± s.e.m. in all figures, unless otherwise noted. Sample sizes (n) denote the number of synapses or cells used as reported in the figure legends or the results. The exact P values and the corresponding inferential statistical methods are stated in the results. Detailed information on statistical tests is provided in Table S1-S11.

