## Supplemental Data 1 for "Systematic microcircuit reconfigurations underlie early experience-induced visual cortical plasticity"

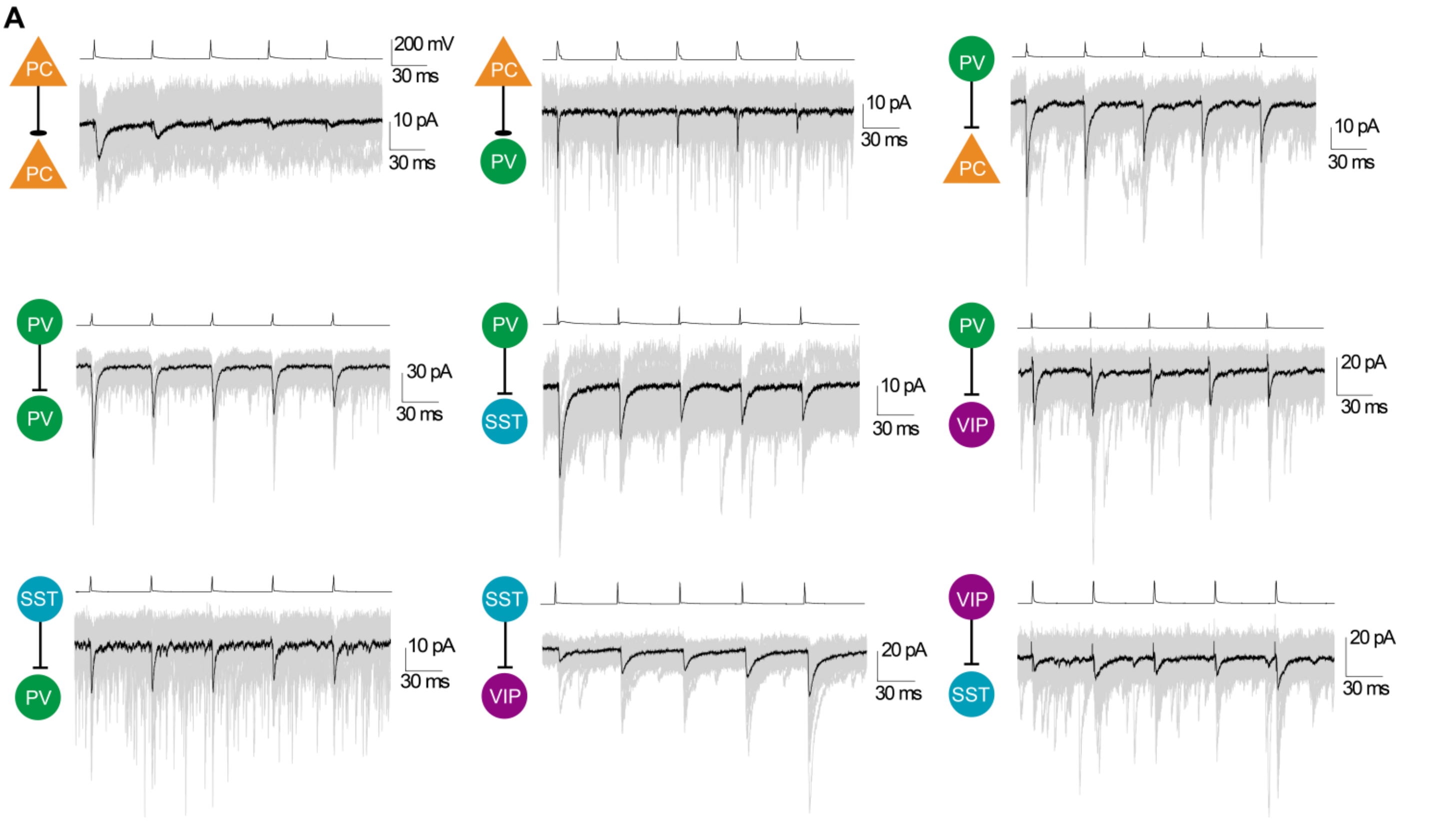


**Figure S1. Representative traces from various synapses in layer 4.** Related to Figure 1.

(A) Representative traces of uEPSCs or uIPSCs recorded from various excitatory, inhibitory or disinhibitory synapses among PCs, PV-, SST- and VIP-INs in the layer 4 microcircuit.

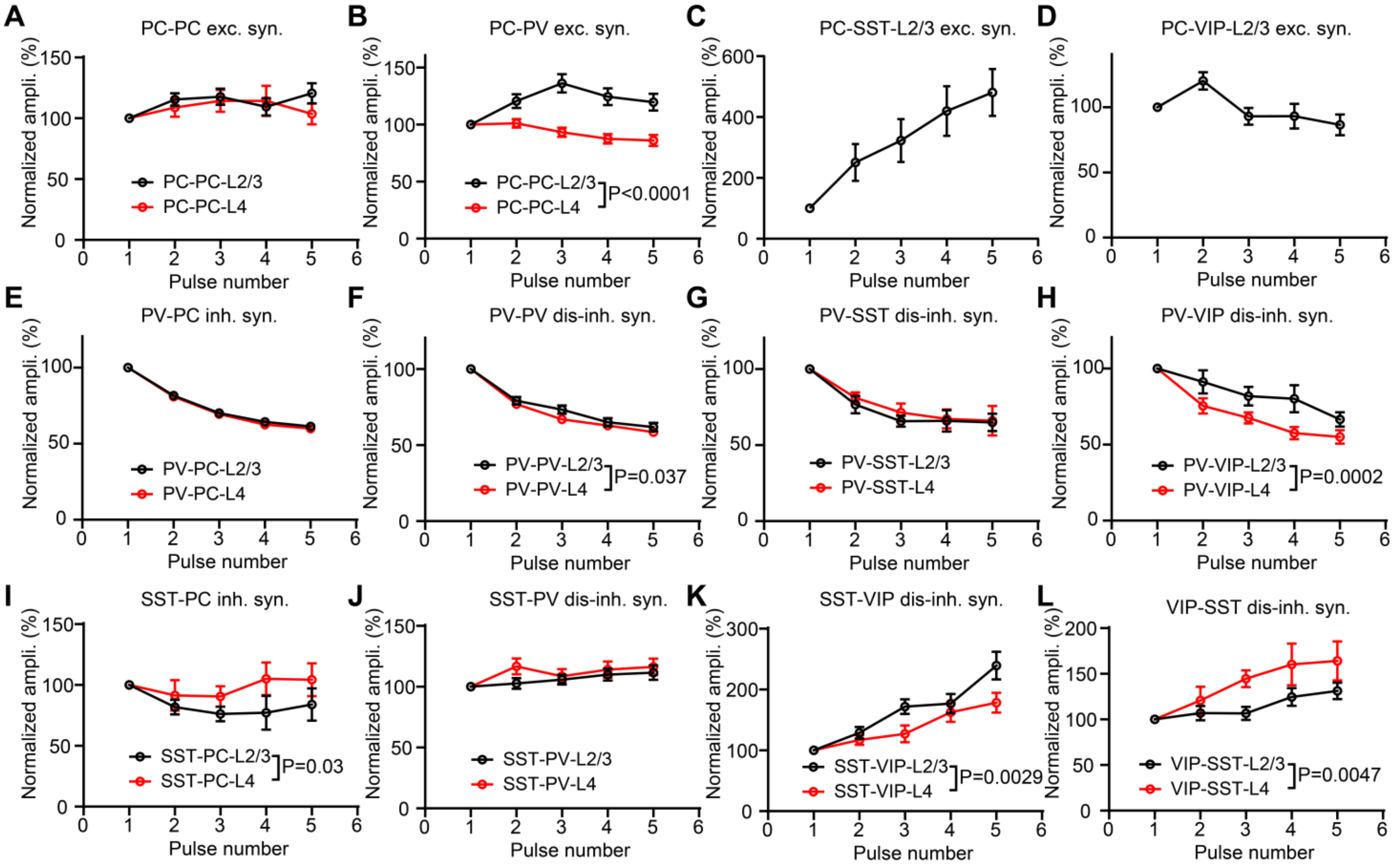


**Figure S2. Short-term synaptic plasticity of various types of intra-cortical synapses.** Related to Figure 1.

(A–L) Depression, facilitation from various synapses in both layers, the synaptic dynamics were tested by 5 presynaptic spikes at 20 Hz for 30 trials.

Data in panels are presented in mean ± s.e.m., the dots represent the connected synapses. See statistical data in Table S6.

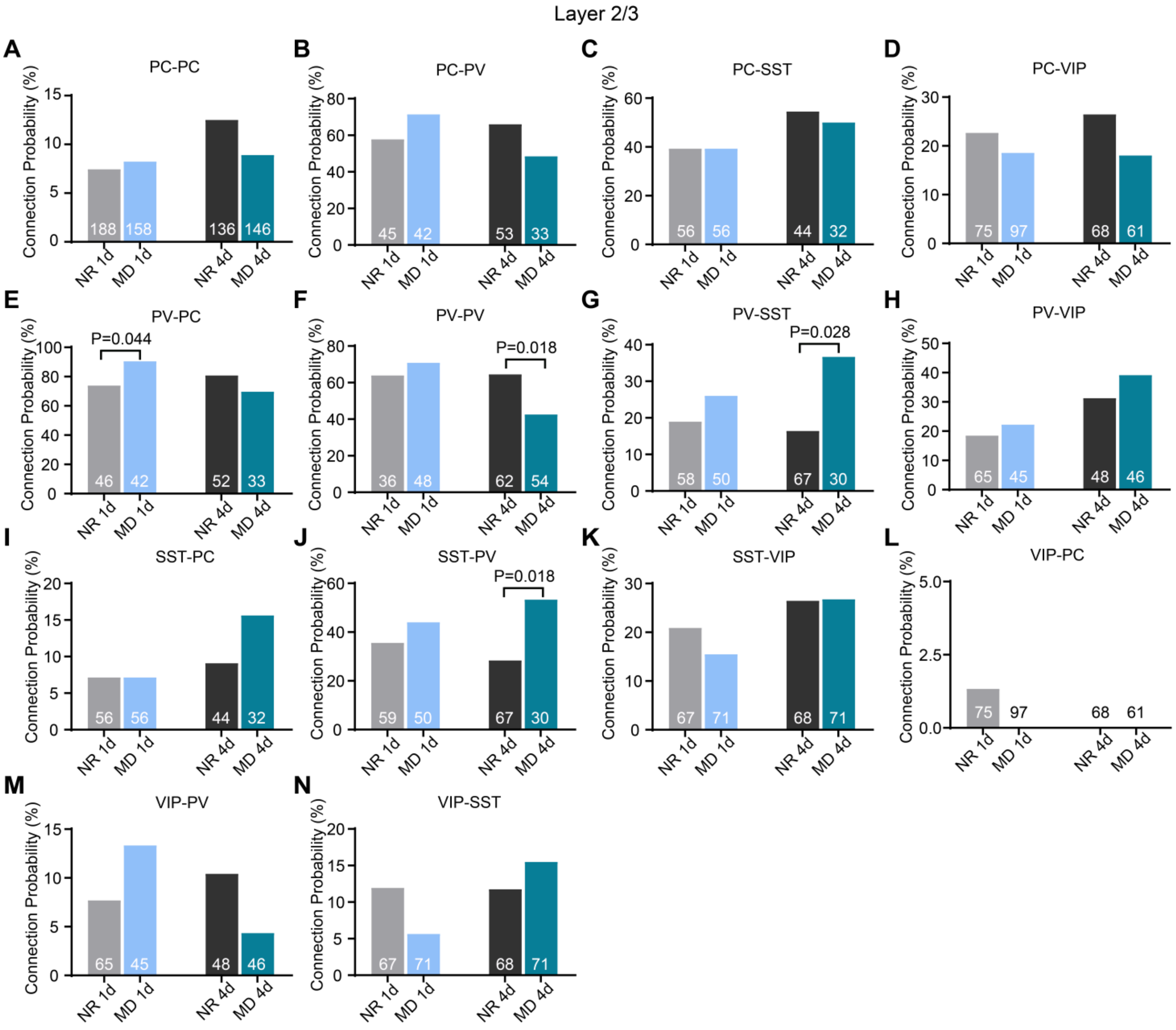


**Figure S3. Connectivity probability from NR and MD groups in layer 2/3.** Related to Figure 2, 3 and 4.

(A–N) Connection probability in pairs from NR 1d, MD 1d and NR 4d, MD 4d groups of different types of synapses in layer 2/3, the number is each group represents total recorded pairs.

Data in panels are presented in mean ± s.e.m., the dots represent the connected synapses. See statistical data in Table S7.

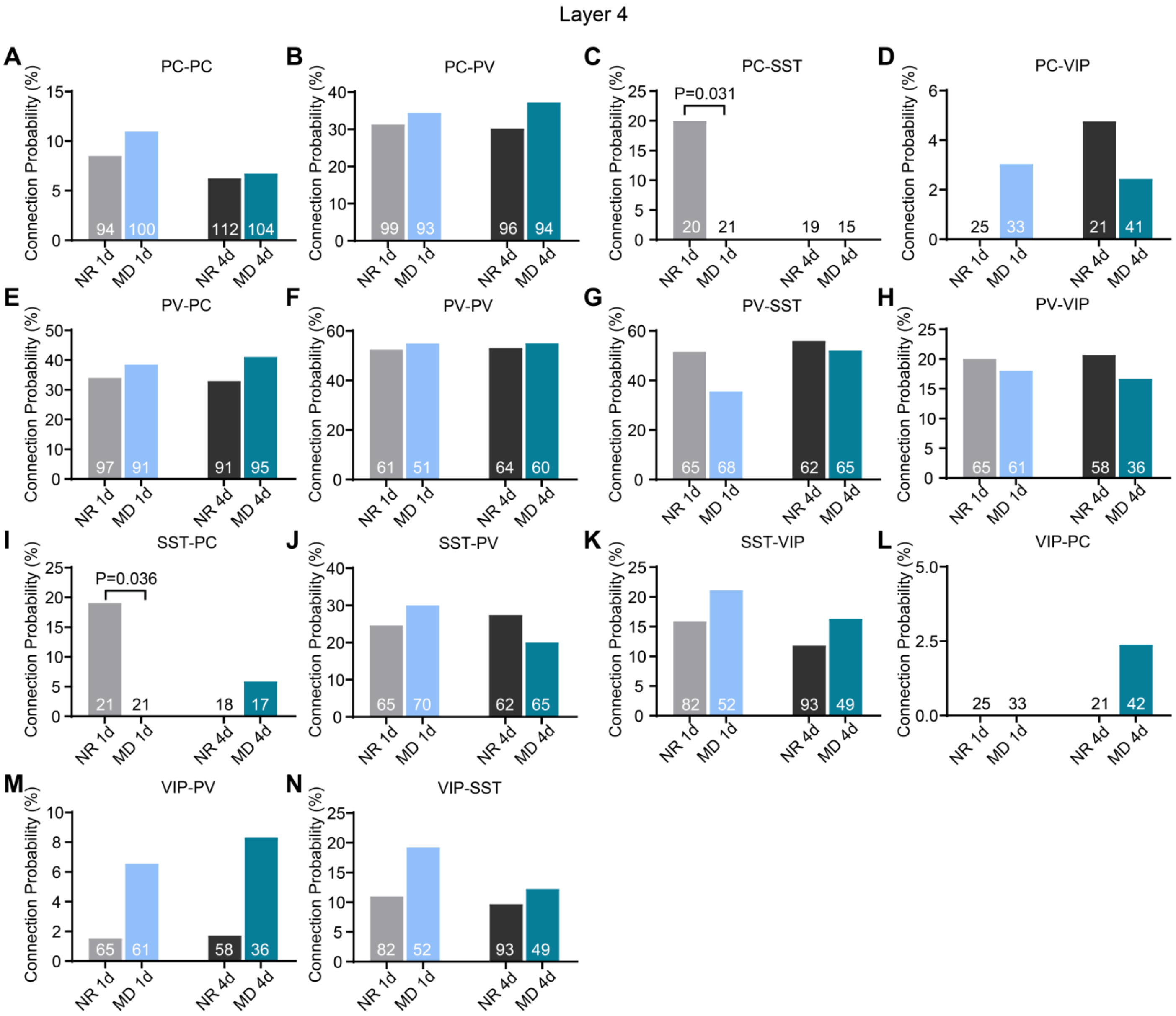


**Figure S4. Connectivity probability from NR and MD groups in layer 4.** Related to Figure 2, 3 and 4.

(A–N) Connection probability in pairs from NR1, MD1 and NR4, MD4 groups of different types of synapses in layer 4, the number is each group represents total recorded pairs.

Data in panels are presented in mean ± s.e.m., the dots represent the connected synapses. See statistical data in Table S8.

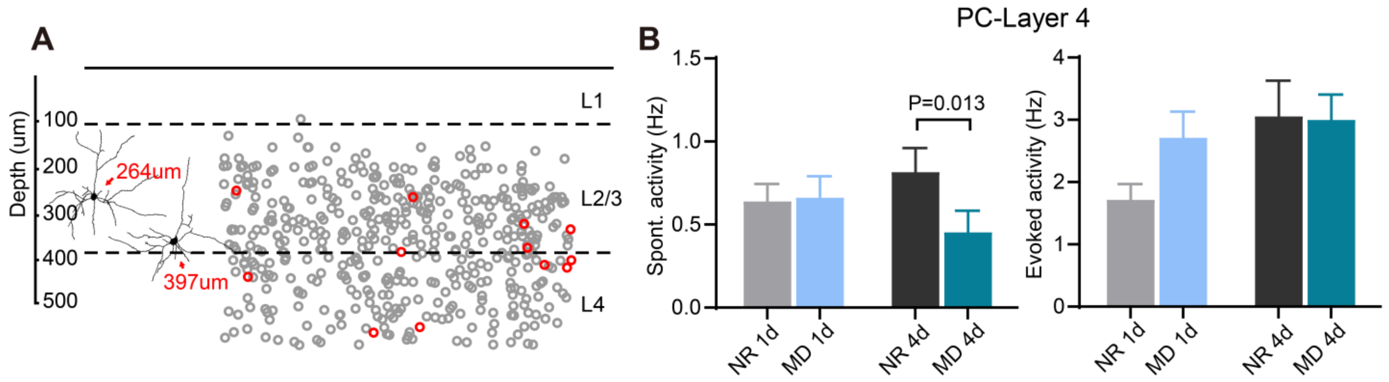


**Figure S5. Neuronal activity changes of PCs in layer 4.** Related to Figure 5.

(A) The distribution of recorded PCs across layers in visual cortex, the morphological reconstructions of neurons represent recorded PCs in layer 2/3.

(B) Changes of spontaneous and evoked activity by 1-day MD and 4-day MD from PCs in layer 4.

Data in panels are presented in mean ± s.e.m., the dots represent the connected synapses. See statistical data in Table S9.

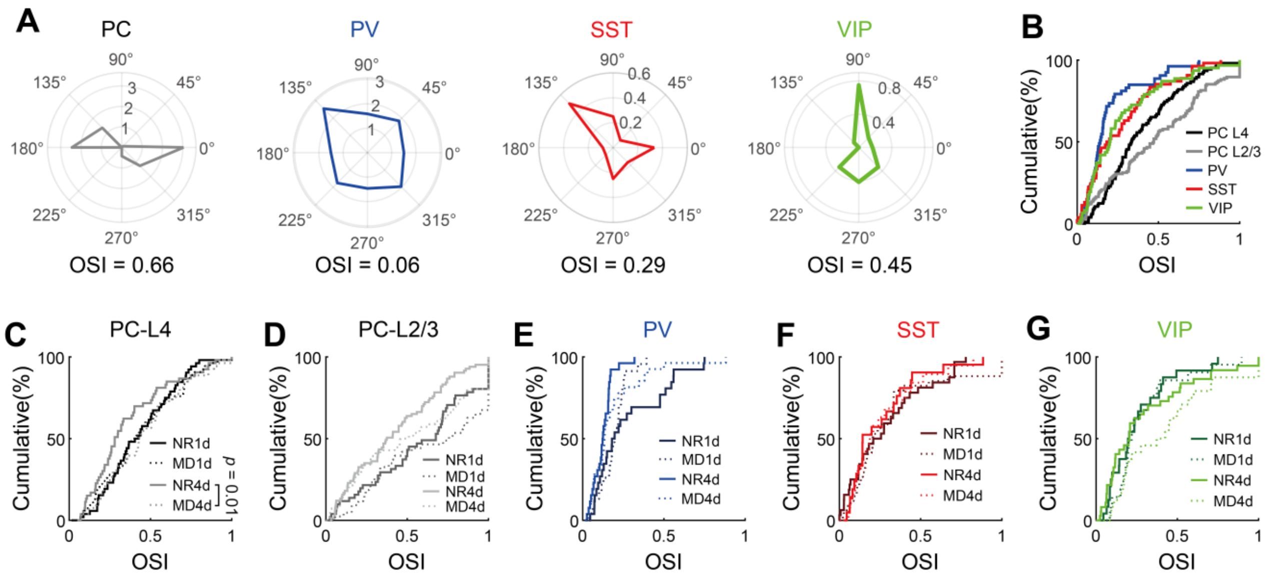


**Figure S6. The orientation selectivity of different subtype neurons.** Related to Figure 5.

(A) Polar plots illustrating orientation selectivity of different subtypes of neurons (from left to right: PC, PV, SST and VIP).

(B) Cumulative distributions of orientation selectivity index (OSI) in normally reared mice. PCs in layer 2/3 showed significantly higher OSI than PV ($p = 8.31\times10^{-10}$), SST ($p = 8.36\times10^{-5}$), and VIP ($p = 4.05\times10^{-6}$) in the same layer. Layer 4 PCs also exhibited higher OSI than L2/3 PV ($p = 8.31\times10^{-10}$), SST ($p = 2.78\times10^{-4}$), and VIP ($p = 1.50\times10^{-4}$). Additionally, layer 2/3 PCs showed higher OSI than L4 PCs ($p = 0.03$).

(C) Cumulative distributions of OSI for layer 4 PCs under 1-day MD, 1-day control, 4-day MD, and 4-day control.

(D–G) Cumulative distributions of OSI for layer 2/3 PCs (D), PV (E), SST (F), and VIP (G), similar to panel (C). (All the p-values are calculated by the Kolmogorov–Smirnov test.)

Data in panels are presented in mean ± s.e.m., the dots represent the connected synapses. See statistical data in Table S10.

**Table S1. Statistics results on synaptic properties and synaptic dynamics of variety synapses within both layers.**

| | | PC-PC | PC-PV | PV-PC | PV-PV | PV-SST | PV-VIP | SST-PV | SST-VIP | VIP-SST |
|---|---|---|---|---|---|---|---|---|---|---|
| Synaptic strength (pA) | L2/3 | 8.1± 1.2 (n = 30) | 22.4 ± 2.3 (n = 41) | 78.9 ± 10.2 (n = 51) | 70.0 ± 12.3 (n = 39) | 41.0 ± 13.8 (n = 21) | 35.9 ± 7.1 (n = 26) | 11.5 ± 1.2 (n = 35) | 6.8 ± 1.4 (n = 25) | 5.2 ± 0.6 (n = 13) |
| | L4 | 5.9±1.2 (n = 15) | 27.1 ± 3.1 (n = 51) | 66.4 ± 11.8 (n = 52) | 95.5 ± 13.8 (n = 56) | 53.4 ± 16.7 (n = 7) | 58.5 ± 10.6 (n = 23) | 19.5 ± 3.3 (n = 28) | 6.9 ± 1.2 (n = 19) | 3.6 ± 10.4 (n = 13) |
| | P-value | 0.218 | 0.2763 | 0.068 | 0.271 | 0.3756 | 0.0987 | 0.063 | 0.5733 | 0.0483 |
| Failure rate (%) | L2/3 | 33.2 ± 5.3 (n = 27) | 21.3 ± 2.3 (n = 41) | 3.8 ± 1.1 (n = 51) | 7.58 ± 3.5 (n = 11) | 12.4 ± 2.8 (n = 21) | 23.9 ± 3.6 (n = 26) | 20.4 ± 2.4 (n = 35) | 56.1 ± 3.9 (n = 25) | 59.6 ± 3.8 (n = 13) |
| | L4 | 57.8 ± 6.7 (n = 14) | 17.4 ± 2.1 (n = 51) | 14.5 ± 2.7 (n = 48) | 12.1 ± 6.1 (n = 14) | 20.5 ± 8.4 (n = 7) | 21.9 ± 4.1 (n = 23) | 24.1 ± 3.7 (n = 28) | 64 ± 3.7 (n = 19) | 61 ± 6.3 (n = 13) |
| | P-value | 0.0086** | 0.1358 | 0.0008*** | 0.706 | 0.5723 | 0.623 | 0.673 | 0.3606 | 0.624 |
| Rise time (ms) | L2/3 | 4.5 ± 1.4 (n = 29) | 0.67 ± 0.07 (n = 41) | 0.59 ± 0.06 (n = 51) | 0.72 ± 0.06 (n = 39) | 0.7 ± 0.09 (n = 21) | 0.84 ± 0.29 (n = 27) | 1.1 ± 0.11 (n = 35) | 0.99 ± 0.11 (n = 25) | 2.03 ± 0.34 (n = 13) |
| | L4 | 1.4 ± 0.3 (n = 15) | 0.55 ± 0.04 (n = 51) | 0.55 ± 0.05 (n = 49) | 0.73 ± 0.05 (n = 56) | 0.54 ± 0.1 (n = 7) | 0.41 ± 0.02 (n = 23) | 0.76 ± 0.07 (n = 28) | 0.71 ± 0.06 (n = 19) | 1.45 ± 0.46 (n = 7) |
| | P-value | 0.0121* | 0.0868 | 0.113 | 0.706 | 0.0706 | 0.1809 | 0.0203* | 0.2204 | 0.3229 |
| Decay time (ms) | L2/3 | 4.0 ± 0.7 (n = 29) | 3.27 ± 0.21 (n = 41) | 8.69 ± 0.33 (n = 51) | 7.02 ± 0.69 (n = 39) | 13.4 ± 1.66 (n = 21) | 7.22 ± 0.54 (n = 27) | 5.02 ± 0.34 (n = 35) | 4.65 ± 0.69 (n = 25) | 10.8 ± 1.97 (n = 9) |
| | L4 | 30. ± 0.47 (n = 15) | 3.31 ± 0.17 (n = 51) | 7.31 ± 0.32 (n = 49) | 5.21 ± 0.16 (n = 56) | 11.6 ± 0.98 (n = 7) | 7.14 ± 0.41 (n = 23) | 4.56 ± 0.3 (n = 28) | 5.34 ± 1.23 (n = 19) | 2.44 ± 0.82 (n = 14) |
| | P-value | 0.7677 | 0.7427 | 0.0021** | 0.0063** | 0.5329 | 0.9111 | 0.3308 | 0.8329 | 0.0003*** |
| Latency (ms) | L2/3 | 1.3 ± 0.07 (n = 27) | 0.86 ± 0.04 (n = 41) | 0.67 ± 0.02 (n = 51) | 0.74 ± 0.05 (n = 11) | 0.8 ± 0.04 (n = 21) | 0.82 ± 0.05 (n = 27) | 0.94 ± 0.03 (n = 35) | 1.0 ± 0.04 (n = 25) | 1.33 ± 0.07 (n = 13) |
| | L4 | 1.18 ± 0.1 (n = 14) | 0.87 ± 0.05 (n = 51) | 0.64 ± 0.02 (n = 48) | 0.79 ± 0.1 (n = 14) | 0.76 ± 0.06 (n = 7) | 0.07 ± 0.04 (n = 23) | 0.86 ± 0.3 (n = 28) | 1.1 ± 0.07 (n = 19) | 1.06 ± 0.06 (n = 12) |
| | P-value | 0.1194 | 0.9049 | 0.503 | 0.8394 | 0.5662 | 0.623 | 0.0598 | 0.192 | 0.0088** |
