## Supplemental Data 2 for "Systematic microcircuit reconfigurations underlie early experience-induced visual cortical plasticity"

**Table S2. Statistics results on the excitatory synapses in layer 2/3 and 4.** It refers to the data shown in Figure 2B–2E, 2G and 2H.

| | Data source | Statistic methods | Statistic results | Synapses number |
|---|---|---|---|---|
| B | PC-PC<br>exc. syn.<br>Layer 2/3<br>NR *vs.* MD | Mann–Whitney<br>*U* test | $U_{1d}$ = 71<br>$P_{1d}$ = 0.3499;<br>$U_{4d}$ = 73<br>$P_{4d}$ = 0.4809 | $NR_{1d}$ = 14 (9 mice)<br>$MD_{1d}$ = 13 (8 mice);<br>$NR_{4d}$ = 16 (7 mice)<br>$MD_{4d}$ = 11 (6 mice) |
| C | PC-PV<br>exc. syn.<br>Layer 2/3<br>NR *vs.* MD | Mann–Whitney<br>*U* test | $U_{1d}$ = 134<br>$P_{1d}$ = 0.0294;<br>$U_{4d}$ = 129<br>$P_{4d}$ = 0.9855 | $NR_{1d}$ = 21 (8 mice)<br>$MD_{1d}$ = 21 (8 mice);<br>$NR_{4d}$ = 20 (6 mice)<br>$MD_{4d}$ = 13 (6 mice) |
| D | PC-SST<br>exc. syn.<br>Layer 2/3<br>NR *vs.* MD | Mann–Whitney<br>*U* test (1d);<br>Unpaired<br>*t* test (4d) | $U_{1d}$ = 91<br>$P_{1d}$ = 0.6471;<br>$t(25)$ = 0.5709<br>$P_{4d}$ = 0.5732 | $NR_{1d}$ = 12 (6 mice)<br>$MD_{1d}$ = 17 (6 mice);<br>$NR_{4d}$ = 15 (5 mice)<br>$MD_{4d}$ = 12 (5 mice) |
| E | PC-VIP<br>exc. syn.<br>Layer 2/3<br>NR *vs.* MD | Mann–Whitney<br>*U* test | $U_{1d}$ = 55<br>$P_{1d}$ = 0.0292;<br>$U_{4d}$ = 49<br>$P_{4d}$ = 0.7544 | $NR_{1d}$ = 15 (9 mice)<br>$MD_{1d}$ = 14 (9 mice);<br>$NR_{4d}$ = 12 (8 mice)<br>$MD_{4d}$ = 9 (7 mice) |
| G | PC-PC<br>exc. syn.<br>Layer 4<br>NR *vs.* MD | Mann–Whitney<br>*U* test | $U_{1d}$ = 37<br>$P_{1d}$ = 0.5999;<br>$U_{4d}$ = 10<br>$P_{4d}$ = 0.0728 | $NR_{1d}$ = 8 (4 mice)<br>$MD_{1d}$ = 11 (4 mice);<br>$NR_{4d}$ = 7 (5 mice)<br>$MD_{4d}$ = 7 (5 mice) |
| H | PC-PV<br>exc. syn.<br>Layer 4<br>NR *vs.* MD | Mann–Whitney<br>*U* test | $U_{1d}$ = 180<br>$P_{1d}$ = 0.002;<br>$U_{4d}$ = 351<br>$P_{4d}$ = 0.8499 | $NR_{1d}$ = 26 (7 mice)<br>$MD_{1d}$ = 27 (7 mice);<br>$NR_{4d}$ = 25 (7 mice)<br>$MD_{4d}$ = 29 (7 mice) |

**Table S3. Statistics results on inhibitory synapses and the Layer 4 input onto PC and PV in layer 2/3.** It refers to the data shown in Figure 3A, 3B, 3D and 3E.

| | Data source | Statistic methods | Statistic results | Neurons number |
|---|---|---|---|---|
| A | PV-PC Inh. syn. Layer 2/3 NR *vs.* MD | Mann–Whitney *U* test | $U_{1d}$ = 199<br>$P_{1d}$ = 0.4898;<br>$U_{4d}$ = 184<br>$P_{4d}$ = 0.0052 | $NR_{1d}$ = 19 (8 mice)<br>$MD_{1d}$ = 24 (8 mice);<br>$NR_{4d}$ = 32 (6 mice)<br>$MD_{4d}$ = 21 (6 mice) |
| B | PV-PC Inh. syn. Layer 4 NR *vs.* MD | Mann–Whitney *U* test | $U_{1d}$ = 321<br>$P_{1d}$ = 0.2627;<br>$U_{4d}$ = 228<br>$P_{4d}$ = 0.1701 | $NR_{1d}$ = 30 (7 mice)<br>$MD_{1d}$ = 26 (7 mice);<br>$NR_{4d}$ = 22 (7 mice)<br>$MD_{4d}$ = 27 (7 mice) |
| D | L4-L2/3-PC 1d NR *vs.* MD | Two-way ANOVA | $F_{interaction\ (5,\ 258)}$ = 0.1444; $P_{interaction}$ = 0.9816<br>$F_{pulse\ (5,\ 258)}$ = 58.10; $P_{pulse}$ <0.0001<br>$F_{1d\ (1,\ 258)}$ = 0.8427; $P_{1d}$ = 0.3595 | $NR_{1d}$ = 22 (3 mice);<br>$MD_{1d}$ = 23 (3 mice) |
| D | L4-L2/3-PC 4d NR *vs.* MD | Two-way ANOVA | $F_{interaction\ (5,\ 246)}$ = 0.2400; $P_{interaction}$ = 0.9444<br>$F_{pulse\ (5,\ 246)}$ = 62.29; $P_{pulse}$ < 0.0001<br>$F_{4d\ (1,\ 246)}$ = 9.421; $P_{4d}$ = 0.0024 | $NR_{4d}$ = 22 (3 mice);<br>$MD_{4d}$ = 21 (3 mice) |
| E | L4-L2/3-PV 1d NR *vs.* MD | Two-way ANOVA | $F_{interaction\ (5,\ 166)}$ = 0.05965; $P_{interaction}$ = 0.9976<br>$F_{pulse\ (5,\ 166)}$ = 37.95; $P_{pulse}$ < 0.0001<br>$F_{1d\ (1,\ 166)}$ = 1.185; $P_{1d}$ = 0.2778 | $NR_{1d}$ = 14 (3 mice);<br>$MD_{1d}$ = 16 (3 mice) |
| E | L4-L2/3-PV 4d NR *vs.* MD | Two-way ANOVA | $F_{interaction\ (5,\ 177)}$ = 0.3011; $P_{interaction}$ = 0.9117<br>$F_{pulse\ (5,\ 177)}$ = 44.15; $P_{pulse}$ < 0.0001<br>$F_{1d\ (1,\ 177)}$ = 2.734; $P_{4d}$ = 0.1000 | $NR_{4d}$ = 17 (3 mice);<br>$MD_{4d}$ = 16 (3 mice) |

**Table S4. Statistics results on the dis-inhibitory synapses in layer 2/3 and 4.** It refers to the data shown in Figure 4B–4G and 4I–4N.

| | Data source | Statistic methods | Statistic results | Synapses number (mice number) |
|---|---|---|---|---|
| B | PV-PV dis-inh. syn. Layer 2/3 NR *vs.* MD | Mann–Whitney *U* test | $U_{1d}$ = 195 $P_{1d}$ = 0.8559; $U_{4d}$ = 191 $P_{4d}$ = 0.2558 | $NR_{1d}$ = 15 (4 mice) $MD_{1d}$ = 27 (4 mice); $NR_{4d}$ = 24 (5 mice) $MD_{4d}$ = 20 (4 mice) |
| C | PV-SST dis-inh. syn. Layer 2/3 NR *vs.* MD | Mann–Whitney *U* test (1d); Unpaired *t* test (4d) | $U_{1d}$ = 53 $P_{1d}$ = 0.3031; $t(17)$ = 0.1629 $P_{4d}$ = 0.8725 | $NR_{1d}$ = 11 (10 mice) $MD_{1d}$ = 13 (10 mice); $NR_{4d}$ = 10 (9 mice) $MD_{4d}$ = 9 (6 mice) |
| D | PV-VIP dis-inh. syn. Layer 2/3 NR *vs.* MD | Mann–Whitney *U* test | $U_{1d}$ = 55 $P_{1d}$ = 0.7713; $U_{4d}$ = 111 $P_{4d}$ = 0.5508 | $NR_{1d}$ = 12 (13 mice) $MD_{1d}$ = 10 (7 mice); $NR_{4d}$ = 15 (9 mice) $MD_{4d}$ = 17 (12 mice) |
| E | SST-PV dis-inh. syn. Layer 2/3 NR *vs.* MD | Mann–Whitney *U* test | $U_{1d}$ = 138 $P_{1d}$ = 0.1565; $U_{4d}$ = 53 $P_{4d}$ = 0.0022 | $NR_{1d}$ = 18 (10 mice) $MD_{1d}$ = 21 (10 mice); $NR_{4d}$ = 17 (9 mice) $MD_{4d}$ = 16 (6 mice) |
| F | SST-VIP dis-inh. syn. Layer 2/3 NR *vs.* MD | Mann–Whitney *U* test | $U_{1d}$ = 35 $P_{1d}$ = 0.1734; $U_{4d}$ = 46 $P_{4d}$ = 0.0091 | $NR_{1d}$ = 10 (8 mice) $MD_{1d}$ = 11 (8 mice); $NR_{4d}$ = 15 (7 mice) $MD_{4d}$ = 14 (9 mice) |
| G | VIP-SST dis-inh. syn. Layer 2/3 NR *vs.* MD | Unpaired *t* test (1d); Mann–Whitney *U* test (4d) | $t(8)$ = 1.885 $P_{1d}$ = 0.0962; $U_{4d}$ = 24 $P_{4d}$ = 0.6943 | $NR_{1d}$ = 6 (7 mice) $MD_{1d}$ = 4 (8 mice); $NR_{4d}$ = 7 (7 mice) $MD_{4d}$ = 8 (7 mice) |
| I | PV-PV dis-inh. syn. Layer 4 NR *vs.* MD | Mann–Whitney *U* test | $U_{1d}$ = 270 $P_{1d}$ = 0.9243; $U_{4d}$ = 344 $P_{4d}$ = 0.1756 | $NR_{1d}$ = 25 (4 mice) $MD_{1d}$ = 22 (4 mice); $NR_{4d}$ = 31 (6 mice) $MD_{4d}$ = 28 (6 mice) |
| J | PV-SST dis-inh. syn. Layer 4 NR *vs.* MD | Unpaired *t* test | $t(6)$ = 0.565 $P_{1d}$ = 0.5925; $t(6)$ = 0.1115 $P_{4d}$ = 0.9149 | $NR_{1d}$ = 4 (7 mice) $MD_{1d}$ = 4 (7 mice); $NR_{4d}$ = 3 (8 mice) $MD_{4d}$ = 5 (7 mice) |
| K | PV-VIP dis-inh. syn. Layer 4 NR *vs.* MD | Mann–Whitney *U* test | $U_{1d}$ = 50 $P_{1d}$ = 0.5190; $U_{4d}$ = 22 $P_{4d}$ = 0.2129 | $NR_{1d}$ = 11 (14 mice) $MD_{1d}$ = 11 (10 mice); $NR_{4d}$ = 12 (13 mice) $MD_{4d}$ = 6 (12 mice) |
| L | SST-PV dis-inh. syn. Layer 4 | Mann–Whitney *U* test (1d); Unpaired | $U_{1d}$ = 131 $P_{1d}$ = 0.9007; $t(23)$ = 2.511 | $NR_{1d}$ = 15 (7 mice) $MD_{1d}$ = 18 (7 mice); $NR_{4d}$ = 13 (8 mice) |

| | | | | |
|---|---|---|---|---|
| | NR *vs.* MD | *t* test (4d) | $P_{4d}$ = 0.0195 | $MD_{4d}$ = 12 (7 mice) |
| M | SST-VIP dis-inh. syn. Layer 4 NR *vs.* MD | Mann–Whitney *U* test (1d); Unpaired *t* test (4d) | $U_{1d}$ = 42 $P_{1d}$ = 0.6027; $t(14)$ = 2.322 $P_{4d}$ = 0.0358 | $NR_{1d}$ = 11 (12 mice) $MD_{1d}$ = 9 (8 mice); $NR_{4d}$ = 8 (14 mice) $MD_{4d}$ = 8 (11 mice) |
| N | VIP-SST dis-inh. syn. Layer 4 NR *vs.* MD | Unpaired *t* test | $t(12)$ = 0.8602 $P_{1d}$ = 0.4066; $t(12)$ = 1.245 $P_{4d}$ = 0.2371 | $NR_{1d}$ = 6 (12 mice) $MD_{1d}$ = 8 (8 mice); $NR_{4d}$ = 9 (14 mice) $MD_{4d}$ = 5 (11 mice) |

**Table S5. Statistics results on the neuronal activity in subtypes of neurons in layer 2/3.** It refers to the data shown in Figure 5C.

| Data source | Statistic methods | Statistic results | Neurons number |
|---|---|---|---|
| PC in L2/3<br>Spont. activity<br>NR *vs.* MD | Mann–Whitney<br>*U* test | $U_{1d} = 345$<br>$P_{1d} = 0.1720$;<br>$U_{4d} = 657$<br>$P_{4d} = 0.0171$ | $NR_{1d} = 29$ (4 mice)<br>$MD_{1d} = 30$ (5 mice);<br>$NR_{4d} = 52$ (6 mice)<br>$MD_{4d} = 36$ (5 mice) |
| PV in L2/3<br>Spont. activity<br>NR *vs.* MD | Mann–Whitney<br>*U* test | $U_{1d} = 231.5$<br>$P_{1d} = 0.8117$;<br>$U_{1d} = 188.5$<br>$P_{1d} = 0.0057$; | $NR_{1d} = 22$ (4 mice)<br>$MD_{1d} = 22$ (3 mice);<br>$NR_{4d} = 25$ (3 mice)<br>$MD_{4d} = 27$ (4 mice) |
| SST in L2/3<br>Spont. activity<br>NR *vs.* MD | Mann–Whitney<br>*U* test | $U_{1d} = 328$<br>$P_{1d} = 0.4251$;<br>$U_{4d} = 142.5$<br>$P_{4d} = 0.9520$ | $NR_{1d} = 25$ (4 mice)<br>$MD_{1d} = 30$ (5 mice);<br>$NR_{4d} = 17$ (3 mice)<br>$MD_{4d} = 17$ (3 mice) |
| VIP in L2/3<br>Spont. activity<br>NR *vs.* MD | Mann–Whitney<br>*U* test | $U_{1d} = 149$<br>$P_{1d} = 0.5085$;<br>$U_{4d} = 300.5$<br>$P_{4d} = 0.0094$ | $NR_{1d} = 18$ (3 mice)<br>$MD_{1d} = 19$ (3 mice);<br>$NR_{4d} = 47$ (5 mice)<br>$MD_{4d} = 21$ (3 mice) |
| PC in L2/3<br>Evoked activity<br>NR *vs.* MD | Mann–Whitney<br>*U* test | $U_{1d} = 372.5$<br>$P_{1d} = 0.3479$;<br>$U_{4d} = 926.5$<br>$P_{4d} = 0.9377$ | $NR_{1d} = 29$ (4 mice)<br>$MD_{1d} = 30$ (5 mice);<br>$NR_{4d} = 52$ (6 mice)<br>$MD_{4d} = 36$ (5 mice) |
| PV in L2/3<br>Evoked activity<br>NR *vs.* MD | Mann–Whitney<br>*U* test | $U_{1d} = 135$<br>$P_{1d} = 0.0114$;<br>$U_{4d} = 335$<br>$P_{4d} = 0.9710$ | $NR_{1d} = 22$ (4 mice)<br>$MD_{1d} = 22$ (3 mice);<br>$NR_{4d} = 25$ (3 mice)<br>$MD_{4d} = 27$ (4 mice) |
| SST in L2/3<br>Evoked activity<br>NR *vs.* MD | Mann–Whitney<br>*U* test | $U_{1d} = 374.5$<br>$P_{1d} = 0.9967$;<br>$U_{4d} = 128$<br>$P_{4d} = 0.5802$ | $NR_{1d} = 25$ (4 mice)<br>$MD_{1d} = 30$ (5 mice);<br>$NR_{4d} = 17$ (3 mice)<br>$MD_{4d} = 17$ (3 mice) |
| VIP in L2/3<br>Evoked activity<br>NR *vs.* MD | Mann–Whitney<br>*U* test | $U_{1d} = 149.5$<br>$P_{1d} = 0.5233$;<br>$U_{4d} = 381$<br>$P_{4d} = 0.1380$ | $NR_{1d} = 18$ (3 mice)<br>$MD_{1d} = 19$ (3 mice);<br>$NR_{4d} = 47$ (5 mice)<br>$MD_{4d} = 21$ (3 mice) |

**Table S6. Statistics results on the short-term plasticity of variety synapses in layers 2/3 and 4.** It refers to the data shown in Figure S2A–S2L.

| | Data source | Statistic methods | Statistic results | Synapses number |
|---|---|---|---|---|
| A | PC-PC exc. syn. L2/3 *vs.* L4 | Two-way ANOVA | $F_{interaction\ (4,\ 215)} = 0.5993$; $P_{interaction} = 0.6636$<br>$F_{pulse\ (4,\ 215)} = 1.316$; $P_{pulse} = 0.2650$<br>$F_{layer\ (1,\ 215)} = 0.8775$; $P_{layer} = 0.3499$ | L2/3 = 30; L4 = 15 |
| B | PC-PV exc. syn. L2/3 *vs.* L4 | Two-way ANOVA | $F_{interaction\ (4,\ 450)} = 5.812$; $P_{interaction} = 1\times10^{-4}$<br>$F_{pulse\ (4,\ 450)} = 2.773$; $P_{pulse} = 0.0268$<br>$F_{layer\ (1,\ 450)} = 69.59$; $P_{layer} < 1\times10^{-4}$ | L2/3 = 41; L4 = 51 |
| E | PV-PC inh. syn. L2/3 *vs.* L4 | Two-way ANOVA | $F_{interaction\ (4,\ 505)} = 0.07916$; $P_{interaction} = 0.9887$<br>$F_{pulse\ (4,\ 505)} = 158.7$; $P_{pulse}<1\times10^{-4}$<br>$F_{layer\ (1,\ 505)} = 0.6590$; $P_{layer} = 0.4173$ | L2/3 = 51; L4 = 52 |
| F | PV-PV inh. syn. L2/3 *vs.* L4 | Two-way ANOVA | $F_{interaction\ (4,\ 465)} = 0.5858$; $P_{interaction} = 0.6731$<br>$F_{pulse\ (4,\ 465)} = 109.3$; $P_{pulse}<1\times10^{-4}$<br>$F_{layer\ (1,\ 465)} = 4.379$; $P_{layer} = 0.0369$ | L2/3 = 39; L4 = 56 |
| G | PV-SST dis-inh. syn. L2/3 *vs.* L4 | Two-way ANOVA | $F_{interaction\ (4,\ 130)} = 0.06575$; $P_{interaction} = 0.9920$<br>$F_{pulse\ (4,\ 130)} = 9.226$; $P_{pulse}<1\times10^{-4}$<br>$F_{layer\ (1,\ 130)} = 0.3503$; $P_{layer} = 0.555$ | L2/3 = 21; L4 = 7 |
| H | PV-VIP dis-inh. syn. L2/3 *vs.* L4 | Two-way ANOVA | $F_{interaction\ (4,\ 240)} = 1.158$; $P_{interaction} = 0.33$<br>$F_{pulse\ (4,\ 240)} = 15.29$; $P_{pulse}<1\times10^{-4}$<br>$F_{layer\ (1,\ 240)} = 13.96$; $P_{layer} = 2\times10^{-4}$ | L2/3 = 27; L4 = 23 |
| I | SST-PC inh. syn. L2/3 *vs.* L4 | Two-way ANOVA | $F_{interaction\ (4,\ 35)} = 0.544$; $P_{interaction} = 0.7042$<br>$F_{pulse\ (4,\ 35)} = 0.822$; $P_{pulse} = 0.5199$<br>$F_{layer\ (1,\ 35)} = 5.101$; $P_{layer} = 0.0302$ | L2/3 = 5; L4 = 4 |
| J | SST-PV dis-inh. syn. L2/3 *vs.* L4 | Two-way ANOVA | $F_{interaction\ (4,\ 305)} = 0.5576$; $P_{interaction} = 0.6936$<br>$F_{pulse\ (4,\ 305)} = 2.336$; $P_{pulse} = 0.0555$<br>$F_{layer\ (1,\ 305)} = 2.593$; $P_{layer} = 0.1084$ | L2/3 = 35; L4 = 28 |
| K | SST-VIP dis-inh. syn. L2/3 *vs.* L4 | Two-way ANOVA | $F_{interaction\ (4,\ 210)} = 1.704$; $P_{interaction} = 0.1502$<br>$F_{pulse\ (4,\ 210)} = 18.56$; $P_{pulse} < 1\times10^{-4}$<br>$F_{layer\ (1,\ 210)} = 9.098$; $P_{\ layer} = 0.0029$ | L2/3 = 25; L4 = 19 |
| L | VIP-SST dis-inh. syn. L2/3 *vs.* L4 | Two-way ANOVA | $F_{interaction\ (4,\ 130)} = 0.7834$; $\mathrm{P}_{interaction} = 0.5379$<br>$F_{pulse\ (4,\ 130)} = 4.476$; $P_{pulse} = 0.002$<br>$F_{layer\ (1,\ 130)} = 8.294$; $P_{layer} = 0.0047$ | L2/3 = 13; L4 = 15 |

**Table S7. Statistics results on the connection probability of variety synapses in layer 2/3.** It refers to the data shown in Figure S3A–S3N.

| | Data source | Statistic methods | Statistic results | Connected synapses/ All synapses |
|---|---|---|---|---|
| A | PC-PC exc. syn. Layer 2/3 NR *vs.* MD | Chi-square test | $P_{1d}$ = 0.7873; $P_{4d}$ = 0.3278 | $NR_{1d}$ = 14/188 $MD_{1d}$ = 13/158; $NR_{4d}$ = 17/136 $MD_{4d}$ = 13/146 |
| B | PC-PV exc. syn. Layer 2/3 NR *vs.* MD | Chi-square test | $P_{1d}$ = 0.1840; $P_{4d}$ = 0.1071 | $NR_{1d}$ = 26/45 $MD_{1d}$ = 30/42; $NR_{4d}$ = 35/53 $MD_{4d}$ = 16/33 |
| C | PC-SST exc. syn. Layer 2/3 NR *vs.* MD | Chi-square test | $P_{1d}$ > 0.9999; $P_{4d}$ = 0.6952 | $NR_{1d}$ = 22/56 $MD_{1d}$ = 22/56; $NR_{4d}$ = 24/44 $MD_{4d}$ = 16/32 |
| D | PC-VIP exc. syn. Layer 2/3 NR *vs.* MD | Chi-square test | $P_{1d}$ = 0.5069; $P_{4d}$ = 0.2517 | $NR_{1d}$ = 17/75 $MD_{1d}$ = 18/97; $NR_{4d}$ = 18/68 $MD_{4d}$ = 11/61 |
| E | PV-PC inh. syn. Layer 2/3 NR *vs.* MD | Chi-square test | $P_{1d}$ = 0.0442; $P_{4d}$ = 0.2409 | $NR_{1d}$ = 34/46 $MD_{1d}$ = 38/42; $NR_{4d}$ = 42/52 $MD_{4d}$ = 23/33 |
| F | PV-PV dis-inh. syn. Layer 2/3 NR *vs.* MD | Chi-square test | $P_{1d}$ = 0.5000; $P_{4d}$ = 0.0181 | $NR_{1d}$ = 23/36 $MD_{1d}$ = 34/48; $NR_{4d}$ = 40/62 $MD_{4d}$ = 23/54 |
| G | PV-SST dis-inh. syn. Layer 2/3 NR *vs.* MD | Chi-square test | $P_{1d}$ = 0.3806; $P_{4d}$ = 0.0277 | $NR_{1d}$ = 11/58 $MD_{1d}$ = 13/50; $NR_{4d}$ = 11/67 $MD_{4d}$ = 11/30 |
| H | PV-VIP dis-inh. syn. Layer 2/3 NR *vs.* MD | Chi-square test | $P_{1d}$ = 0.6278; $P_{4d}$ = 0.4236 | $NR_{1d}$ = 12/65 $MD_{1d}$ = 10/45; $NR_{4d}$ = 15/48 $MD_{4d}$ = 18/46 |
| I | SST-PC inh. syn. Layer 2/3 NR *vs.* MD | Chi-square test | $P_{1d}$ > 0.9999; $P_{4d}$ = 0.2168 | $NR_{1d}$ = 4/56 $MD_{1d}$ = 4/56; $NR_{4d}$ = 4/44 $MD_{4d}$ = 5/32 |
| J | SST-PV dis-inh. syn. Layer 2/3 | Chi-square test | $P_{1d}$ = 0.3709; $P_{4d}$ = 0.0179 | $NR_{1d}$ = 21/59 $MD_{1d}$ = 22/50; $NR_{4d}$ = 19/67 |

| | | | | |
|---|---|---|---|---|
| | NR *vs*. MD | | | $MD_{4d}$ = 16/30 |
| K | SST-VIP<br>dis-inh. syn.<br>Layer 2/3<br>NR *vs*. MD | Chi-square<br>test | $P_{1d}$ = 0.4102;<br>$P_{4d}$ = 0.6836 | $NR_{1d}$ = 14/67<br>$MD_{1d}$ = 11/71;<br>$NR_{4d}$ = 18/68<br>$MD_{4d}$ = 19/71 |
| L | VIP-PC<br>inh. syn.<br>Layer 2/3<br>NR *vs*. MD | Chi-square<br>test | $P_{1d}$ = 0.2540; | $NR_{1d}$ = 1/75<br>$MD_{1d}$ = 0/97;<br>$NR_{4d}$ = 0/68<br>$MD_{4d}$ = 0/61 |
| M | VIP-PV<br>dis-inh. syn.<br>Layer 2/3<br>NR *vs*. MD | Chi-square<br>test | $P_{1d}$ = 0.3322;<br>$P_{4d}$ = 0.2626 | $NR_{1d}$ = 5/65<br>$MD_{1d}$ = 6/45;<br>$NR_{4d}$ = 5/48<br>$MD_{4d}$ = 2/46 |
| N | VIP-SST<br>dis-inh. syn.<br>Layer 2/3<br>NR *vs*. MD | Chi-square<br>test | $P_{1d}$ = 0.1888;<br>$P_{4d}$ = 0.5224 | $NR_{1d}$ = 8/67<br>$MD_{1d}$ = 4/71;<br>$NR_{4d}$ = 8/68<br>$MD_{4d}$ = 11/71 |

**Table S8. Statistics results on the connection probability of variety synapses in layer 4.** It refers to the data shown in Figure S4A–S4N.

| | Data source | Statistic methods | Statistic results | Connected synapses/ All synapses |
|---|---|---|---|---|
| A | PC-PC exc. syn. Layer 4 NR *vs.* MD | Chi-square test | $P_{1d}$ = 0.5599; $P_{4d}$ = 0.8860 | $NR_{1d}$ = 8/94 $MD_{1d}$ = 11/100; $NR_{4d}$ = 7/112 $MD_{4d}$ = 7/104 |
| B | PC-PV exc. syn. Layer 4 NR *vs.* MD | Chi-square test | $P_{1d}$ = 0.6480; $P_{4d}$ = 0.3056 | $NR_{1d}$ = 31/99 $MD_{1d}$ = 32/93; $NR_{4d}$ = 29/96 $MD_{4d}$ = 35/94 |
| C | PC-SST exc. syn. Layer 4 NR *vs.* MD | Chi-square test | $P_{1d}$ = 0.031 | $NR_{1d}$ = 4/20 $MD_{1d}$ = 0/21; $NR_{4d}$ = 0/19 $MD_{4d}$ = 0/15 |
| D | PC-VIP exc. syn. Layer 4 NR *vs.* MD | Chi-square test | $P_{1d}$ = 0.3799; $P_{4d}$ = 0.6242 | $NR_{1d}$ = 0/25 $MD_{1d}$ = 1/33; $NR_{4d}$ = 1/21 $MD_{4d}$ = 1/41 |
| E | PV-PC inh. syn. Layer 4 NR *vs.* MD | Chi-square test | $P_{1d}$ = 0.5265; $P_{4d}$ = 0.2538 | $NR_{1d}$ = 33/97 $MD_{1d}$ = 35/91; $NR_{4d}$ = 30/91 $MD_{4d}$ = 39/95 |
| F | PV-PV dis-inh. syn. Layer 4 NR *vs.* MD | Chi-square test | $P_{1d}$ = 0.7963; $P_{4d}$ = 0.8342 | $NR_{1d}$ = 32/61 $MD_{1d}$ = 28/51; $NR_{4d}$ = 34/64 $MD_{4d}$ = 33/60 |
| G | PV-SST dis-inh. syn. Layer 4 NR *vs.* MD | Chi-square test | $P_{1d}$ = 0.1961; $P_{4d}$ = 0.0854 | $NR_{1d}$ = 8/65 $MD_{1d}$ = 4/68; $NR_{4d}$ = 5/62 $MD_{4d}$ = 12/65 |
| H | PV-VIP dis-inh. syn. Layer 4 NR vs. MD | Chi-square test | $P_{1d}$ = 0.7787; $P_{4d}$ = 0.6299 | $NR_{1d}$ = 13/65 $MD_{1d}$ = 11/61; $NR_{4d}$ = 12/58 $MD_{4d}$ = 6/36 |
| I | SST-PC inh. syn. Layer 4 NR vs. MD | Chi-square test | $P_{1d}$ = 0.0355; $P_{4d}$ = 0.2965 | $NR_{1d}$ = 4/21 $MD_{1d}$ = 0/21; $NR_{4d}$ = 0/18 $MD_{4d}$ = 1/17 |
| J | SST-PV dis-inh. syn. Layer 4 | Chi-square test | $P_{1d}$ = 0.4834; $P_{4d}$ = 0.3251 | $NR_{1d}$ = 16/65 $MD_{1d}$ = 21/70; $NR_{4d}$ = 17/62 |

| | | | | |
|---|---|---|---|---|
| | NR vs. MD | | | $MD_{4d}$ = 13/65 |
| K | SST-VIP<br>dis-inh. syn.<br>Layer 4<br>NR vs. MD | Chi-square<br>test | $P_{1d}$ = 0.4355;<br>$P_{4d}$ = 0.4541 | $NR_{1d}$ = 13/82<br>$MD_{1d}$ = 11/52;<br>$NR_{4d}$ = 11/93<br>$MD_{4d}$ = 8/49 |
| L | VIP-PC<br>inh. syn.<br>Layer 4<br>NR vs. MD | Chi-square<br>test | $P_{4d}$ = 0.4760; | $NR_{1d}$ = 0/25<br>$MD_{1d}$ = 0/33;<br>$NR_{4d}$ = 0/21<br>$MD_{4d}$ = 1/42 |
| M | VIP-PV<br>dis-inh. syn.<br>Layer 4<br>NR vs. MD | Chi-square<br>test | $P_{1d}$ = 0.1492;<br>$P_{4d}$ = 0.1228 | $NR_{1d}$ = 1/65<br>$MD_{1d}$ = 4/61;<br>$NR_{4d}$ = 1/58<br>$MD_{4d}$ = 3/36 |
| N | VIP-SST<br>dis-inh. syn.<br>Layer 4<br>NR vs. MD | Chi-square<br>test | $P_{1d}$ = 0.1819;<br>$P_{4d}$ = 0.6361 | $NR_{1d}$ = 9/82<br>$MD_{1d}$ = 10/52;<br>$NR_{4d}$ = 9/93<br>$MD_{4d}$ = 6/49 |

**Table S9. Statistics results on the neuronal activity of PCs in layer 4.** It refers to the data shown in Figure S5B.

| Data source | Statistic methods | Statistic results | Neurons number |
|---|---|---|---|
| PC in L4<br>Spont. activity<br>NR *vs.* MD | Mann–Whitney<br>*U* test | $U_{1d}$ = 498.5<br>$P_{1d}$ = 0.7029;<br>$U_{4d}$ = 416<br>$P_{4d}$ = 0.0133 | $NR_{1d}$ = 32 (4 mice)<br>$MD_{1d}$ = 33 (4 mice);<br>$NR_{4d}$ = 35 (5 mice)<br>$MD_{4d}$ = 36 (5 mice) |
| PC in L4<br>Evoked activity<br>NR *vs.* MD | Mann–Whitney<br>*U* test | $U_{1d}$ = 393.5<br>$P_{1d}$ = 0.0781;<br>$U_{1d}$ = 546<br>$P_{1d}$ = 0.339; | $NR_{1d}$ = 32 (4 mice)<br>$MD_{1d}$ = 33 (4 mice);<br>$NR_{4d}$ = 35 (5 mice)<br>$MD_{4d}$ = 36 (5 mice) |

**Table S10. Statistical results on the orientation selectivity in subtypes of neurons.** It refers to the data shown in Figure S6B–S6G.

| Data source | Statistic methods | Statistic results | Neurons number |
|---|---|---|---|
| PC, PV, SST, VIP in L2/3 and PC in L4 OSI NR | Kolmogorov–Smirnov test | $P_{(PV, PC L4)}$ = 3.92×10$^{-9}$; $P_{(PV, PC L2/3)}$ = 8.31×10$^{-10}$; $P_{(PV, SST)}$ = 0.0537; $P_{(PV, VIP)}$ = 0.0178; $P_{(PC L4, PC L2/3)}$ = 0.0266; $P_{(SST, PC L4)}$ = 0.0003; $P_{(VIP, PC L4)}$ = 0.0002; $P_{(SST, PC L2/3)}$ = 8.36×10$^{-5}$; $P_{(VIP, PC L2/3)}$ = 4.05×10$^{-6}$; $P_{(SST, VIP)}$ = 0.9656 | $PC_{L4}$ = 107 (9 mice); $PC_{L2/3}$ = 136 (10 mice); PV = 54 (7 mice); SST = 55 (7 mice); VIP = 63 (8 mice) |
| PC in L2/3 OSI NR *vs*. MD | Kolmogorov–Smirnov test | $P_{1d}$ = 0.1595; $P_{4d}$ = 0.0627 | $NR_{1d}$ = 52 (4 mice) $MD_{1d}$ = 44 (5 mice); $NR_{4d}$ = 84 (6 mice) $MD_{4d}$ = 57 (5 mice) |
| PC in L4 OSI NR *vs*. MD | Kolmogorov–Smirnov test | $P_{1d}$ = 0.9042; $P_{4d}$ = 0.0135 | $NR_{1d}$ = 53 (4 mice) $MD_{1d}$ = 55 (4 mice); $NR_{4d}$ = 54 (5 mice) $MD_{4d}$ = 54 (5 mice) |
| PV in L2/3 OSI NR *vs*. MD | Kolmogorov–Smirnov test | $P_{1d}$ = 0.1771; $P_{4d}$ = 0.0057 | $NR_{1d}$ = 27 (4 mice) $MD_{1d}$ = 24 (3 mice); $NR_{4d}$ = 27 (3 mice) $MD_{4d}$ = 28 (4 mice) |
| SST in L2/3 OSI NR *vs*. MD | Kolmogorov–Smirnov test | $P_{1d}$ = 0.9024; $P_{4d}$ = 0.7720 | $NR_{1d}$ = 33 (4 mice) $MD_{1d}$ = 35 (5 mice); $NR_{4d}$ = 22 (3 mice) $MD_{4d}$ = 20 (3 mice) |
| VIP in L2/3 OSI NR *vs*. MD | Kolmogorov–Smirnov test | $P_{1d}$ = 0.5777; $P_{4d}$ = 0.2180 | $NR_{1d}$ = 25 (3 mice) $MD_{1d}$ = 22 (3 mice); $NR_{4d}$ = 38 (5 mice) $MD_{4d}$ = 25 (3 mice) |

OSI: orientation selectivity index

**Table S11. Parameter setting in the microcircuit model.** It refers to the data shown in Figure 6 and 7.

| Parameter | Value |
|---|---|
| **cell-related parameters** | |
| #PC | 700 |
| #PV | 100 |
| #SST | 100 |
| #VIP | 100 |
| time constant $\tau_{PC}$ | 10 ms |
| time constant $\tau_{PV}$ | 10 ms |
| time constant $\tau_{SST}$ | 10 ms |
| time constant $\tau_{VIP}$ | 10 ms |
| rheobase $\theta$ | 14 |
| proportion of currents leaking away from the soma $\lambda_S$ | 0.31 |
| proportion of currents leaking away from the dendrites $\lambda_D$ | 0.27 |
| dendritic signal threshold $\theta_c$ | 28 |
| spike gain factor $c$ | 7 |
| **network input under spontaneous activity** | |
| $x_S$ for MD 4d / otherwise | 17.4 / 18 |
| $x_D$ | 5 |
| $x_{PV}$ | 3.1 |
| $x_{SST}$ | 1.9 |
| $x_{VIP}$ | 1.4 |
| **network input under evoked activity** | |
| $x_S$ | 20.7 / 22.8 |
| $x_D$ | 10 |
| $x_{PV}$ | 7.1 |
| $x_{SST}$ | 3.3 |
| $x_{VIP}$ | 2.8 |
| **Simulation parameters** | |
| time step $\delta t$ | 0.1 ms |
| simulation time | 1000 ms |
